# Identifying simultaneous inhibition of riboflavin metabolism and MEK as a novel therapeutic strategy for pancreatic cancer

**DOI:** 10.1101/2024.10.23.619807

**Authors:** Takako Ooshio, Yusuke Satoh, Hui Jiang, Tongwei Liu, Kiyonaga Fujii, Takamasa Ishikawa, Kaori Igarashi, Kaori Saitoh, Keiko Kato, Tetsuo Mashima, Shigeki Jin, Kotaro Matoba, Matthias Mack, Tsuyoshi Osawa, Hiroyuki Seimiya, Tomoyoshi Soga, Masahiro Sonoshita

**Author notes:** Corresponding authors: Takako Ooshio, PhD., Masahiro Sonoshita, PhD.

## Abstract

Pancreatic ductal adenocarcinoma (PDAC) remains a formidable challenge with a survival rate of approximately 10%, largely due to the lack of diagnostic and therapeutic options. To identify novel therapeutic targets for PDAC, we previously developed a *Drosophila melanogaster* model, the ’*4-hit*’ fly, which mimics the alterations in *KRAS*, *TP53*, *CDKN2A*, and *SMAD4* genes observed in PDAC. Through whole-body genetic screening using the *4-hit* model, we identified riboflavin (RF) metabolism as a novel potential therapeutic target for PDAC. Additionally, transcript levels of four out of five genes that are involved in RF metabolism were elevated in human PDAC compared to non-tumor tissues. Roseoflavin (RoF), an antimetabolite of RF, resulted in a variety range of metabolic changes, decreased mitochondrial respiratory chain activity, and suppressed proliferation in cultured PDAC cells. Furthermore, the combination of RoF with the MEK inhibitor drug trametinib significantly suppressed the growth of human PDAC xenografts compared to either treatment alone. These findings underscore that dual targeting of RF metabolism and MEK represents a novel therapeutic strategy for PDAC.

## Introduction

Pancreatic ductal adenocarcinoma (PDAC) remains a formidable challenge due to its late detection and limited effective treatments (Siegel et al, 2023). With a 5-year survival rate of only about 10% in the USA, its prognosis is dire (Mizrahi et al, 2020). The global incidence and mortality rates among PDAC patients are projected to continue rising in the upcoming years (Rahib et al, 2021). PDAC is characterized by frequent alterations such as activation of the oncogene *KRAS* and inactivation of the tumor suppressor genes including *CDKN2A*, *TP53*, and *SMAD4* (Morris et al, 2010). Consequently, patients harboring alterations in all of these genes (hereafter referred to as ‘*4-hit*’) demonstrate the worst prognosis among PDAC patients (Qian et al, 2018). Despite this, there have been no reports of mouse models that develop endogenous *4-hit* tumors, thus hindering the development of novel treatments for *4-hit* PDAC.

To address this issue, we explored the use of *Drosophila melanogaster* as a novel animal model to mimic various PDAC genotypes, leveraging our expertise with mammalian assays for studying gastrointestinal cancers (Sonoshita et al, 2001; Sonoshita et al, 2002; Sonoshita et al, 2011; Sonoshita et al 2015; Itatani et al, 2016; Kakizaki et al, 2017; Okada et al, 2017). Due to genetic similarities to humans and the ease of genetic manipulation, *Drosophila* has proven highly suitable for cancer research (Pandey and Nichols, 2011). For example, the generation of gene knockdown and knockout strains is relatively easy, allowing for comprehensive genetic screening at a whole-body level (Yamamura et al, 2021). Leveraging these capabilities, a *Ret*^M955T^ fly model was developed to mimic the *RET*^M918T^ gene mutation, commonly observed in medullary thyroid carcinoma (MTC), leading to the successful validation of the tyrosine kinase inhibitor drug vandetanib as the first targeted therapy to for the treatment of MTC (Vidal et al, 2005). Furthermore, we devised a novel strategy to reduce the toxicity of the approved kinase inhibitor sorafenib by identifying its ‘anti-targets’, whose inhibition contributes to the drug’s adverse effects (Sonoshita and Cagan, 2017; Sonoshita et al, 2017; Ung et al, 2019). Building on these efforts, we have recently established a *Drosophila* model that replicates the *4-hit* signature (Sekiya et al, 2023). Specifically, we targeted the cDNA sequences of *Drosophila Ras85D* (an ortholog of human *KRAS*) with G12D missense mutation (*dRas*^G12D^) and *Cyclin E* (*dCycE*), along with shRNA sequences targeting the *Drosophila* orthologs of *TP53* and *SMAD4*, using the *Serrate* (*Ser*) enhancer to induce their expression in the wing discs of larvae (Sekiya et al, 2023).

Our studies utilizing these *Ser-gal4,UAS-GFP,UAS-Ras*^G12D^*,UAS-dCycE,UAS- p53*^shRNA^*,UAS-Med*^shRNA^ flies (hereafter referred to as *Ser*>*4-hit* or *4-hit* flies) have demonstrated the utility of the wing disc as a valuable tissue for studying epithelial transformation characterized by increased cell proliferation and migration (Sekiya et al, 2023). Notably, a large portion of *4-hit* flies exhibited lethality. Our genetic screening of the entire kinome revealed that reducing a copy of *Mitogen-activated protein kinase kinase* (*MEK*), *Aurora kinase B* (*AURKB*), and *Glycogen synthase kinase 3* (*GSK3*), enhanced the viability of *4-hit* flies, suggesting that their products are therapeutic targets in PDAC (Sekiya et al., 2023; Fukuda et al., 2024). Furthermore, chemical screenings using *4-hit* flies led to the identification of novel therapeutic candidates that were effective in suppressing xenograft growth of human PDAC in mice, such as a combination of the MEK inhibitor drug trametinib with an AURKB inhibitor BI-8312366 or a GSK3 inhibitor CHIR99021 (Sekiya et al, 2023; Fukuda et al, 2024). Consequently, our approach of utilizing *4-hit* flies in combination with mammalian models provides a promising platform for identifying novel therapeutic targets and candidates for PDAC.

Building upon these achievements, this paper focuses on a Riboflavin kinase (RFK; also called flavokinase, EC 2.7.1.26), a key enzyme in riboflavin (RF) metabolism, ubiquitous across all organisms (Barile et al, 2016). RF, commonly known as vitamin B2, is an essential water-soluble nutrient primarily obtained through dietary intake (Olfat et al, 2022). While excess RF is safely excreted in urine, RF deficiency can lead to a range of clinical symptoms, including cheilosis, angular stomatitis, glossitis, seborrheic dermatitis, and severe anemia with erythroid hypoplasia (McNulty et al, 2023). Epidemiological studies have associated RF deficiency with an increased risk of cancer development (Powers, 2003; Saedisomeolia and Ashoori, 2018). Conversely, other studies suggest that RF metabolism promotes cisplatin resistance in prostate and ovarian cancers (Hirano et al, 2011; Yang et al, 2019). Despite these reported connections, a role of RF metabolism in PDAC pathogenesis remains uninvestigated. Here, we propose the concurrent inhibition of RF metabolism and MEK as a novel therapeutic strategy for PDAC.

## RESULTS

### *Drosophila* genetic screening identifies riboflavin metabolism as a potential therapeutic target for PDAC

We have successfully identified several potential therapeutic targets for PDAC by comprehensive kinome screening utilizing *4-hit* flies (Sekiya et al, 2023; Fukuda et al, 2024). Our library of kinase-mutant flies included not only protein kinases and lipid kinases but also vitamin kinases. Interestingly, we discovered that genetic inhibition of one of the vitamin kinases, *Riboflavin kinase* (*RFK*), significantly rescued *4-hit* fly lethality (Fig. 1A). RFK phosphorylates RF taken up by RF transporters (RFTs) from extracellular space, functioning as the rate-limiting enzyme in RF metabolism (Fig. 1B; Barile et al, 2016; Pedrolli et al, 2011). Once in the cytoplasm flavin mononucleotide (FMN) is synthesized by the RFK from RF and ATP. FAD synthetase 1 (FLAD1) subsequently generates flavin adenine dinucleotide (FAD) from FMN and ATP. Flavoproteins, which depend on FMN and FAD, participate in a wide range of cellular processes, such as energy metabolism and antioxidant responses (Fig. 1B; Lienhart et al, 2013), yet the specific roles of RF metabolism in PDAC pathogenesis had to be previously elucidated.

**Figure 1.**
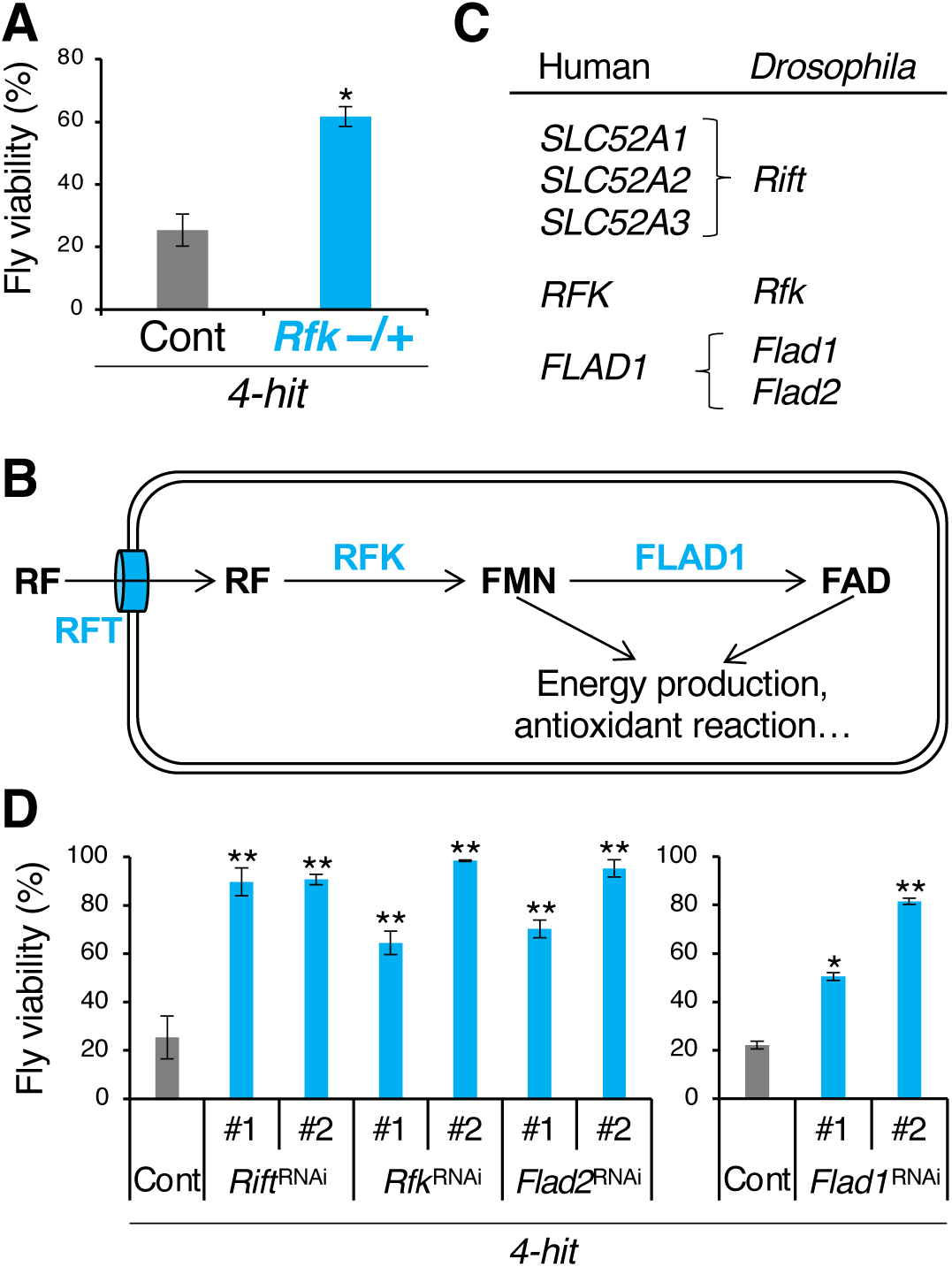
*Drosophila* screening utilizing *4-hit* flies identifies riboflavin metabolism as a potential therapeutic target for PDAC. (A) A heterozygous mutation of *Riboflavin kinase* (*RFK*) rescued *4-hit* fly lethality. (B) Riboflavin (RF) metabolism. RFT, RF transporter. FMN, flavin mononucleotide. FAD, flavin adenine dinucleotide. FLAD1, FAD synthetase 1. (C) RF metabolism-related genes between humans and *Drosophila*. *Solute carrier family 52 member 1* (*SLC52A1*), *SLC52A2*, and *SLC52A3*: *RFTs*. (D) Knockdown of *Rift*, *Rfk*, *Flad1*, or *Flad2* in transformed cells rescued *4-hit* fly lethality. Two RNAi strains (#1 and #2) were tested for each gene. Error bars, standard deviation (SD) in technical triplicate. *, *P* < 0.05; **, *P* < 0.01 in Dunnett’s test compared to control *4-hit* flies.

According to DIOPT ortholog finder, *Drosophila* possesses orthologs of human *RFT*, *RFK*, and *FLAD1* genes, designated as *Rift*, *Rfk*, and *Flad1* and *Flad2*, respectively (Fig. 1C). To further assess the impact of disrupting RF metabolism on *4-hit* fly lethality, we knocked down these genes specifically in transformed cells in the model. Consequently, these knockdown manipulations also rescued the lethality of the *4-hit* flies (Fig. 1D). Furthermore, we examined the expression of genes involved in RF metabolism in PDAC using the GEPIA2 database. Humans possess three *RFT* genes—*Solute carrier family 52 member A1* (*SLC52A1*), *SLC52A2*, *SLC52A3*—as well as *RFK* and *FLAD1*. The expression levels of RF metabolism-related genes with the exception of *SLC52A1*, were elevated in human PDAC tissues compared to non-tumor pancreatic tissues (Fig. EV1). These findings suggest that RF metabolism could serve as a novel therapeutic target for PDAC.

### Long-term roseoflavin treatment suppresses PDAC cell proliferation and mitochondrial respiratory chain activity by reducing RF metabolites in the cytosol

To elucidate the role of RF metabolism in human PDAC cells, we investigated the effects of RF inhibitors on AsPC-1 cells harboring the *4-hit* signature. Among specific inhibitors tested, we particularly focused on roseoflavin (RoF), because it effectively inhibited RF metabolism. RoF, a naturally occurring analog of RF, is produced by the bacteria *Streptomyces davaonensis*, *Streptomyces cinnabarinus* and *Streptomyces berlinensis* (Otani et al, 1980; Landwehr et al, 2018; Liunardo et al, 2024; Jankowitsch et al, 2012). Noted for its competitive inhibition of RF metabolism and its antibiotic properties, RoF has been extensively studied in bacterial systems (Mansjo et al, 2011; Mack and Grill, 2006; Pedrolli et al, 2014; Pedrolli et al, 2012). The mechanism of action involves RoF being transported into the cytosol via RFT, where it is modified by RFK and FLAD1 to form RoF mononucleotide (RoFMN) and RoF adenine dinucleotide (RoFAD), in a manner analogous to RF (Fig. 2A; Pedrolli et al, 2011). Despite its well-documented effects on bacteria, the impacts of RoF on mammalian cells have remained largely unexplored.

**Figure 2.**
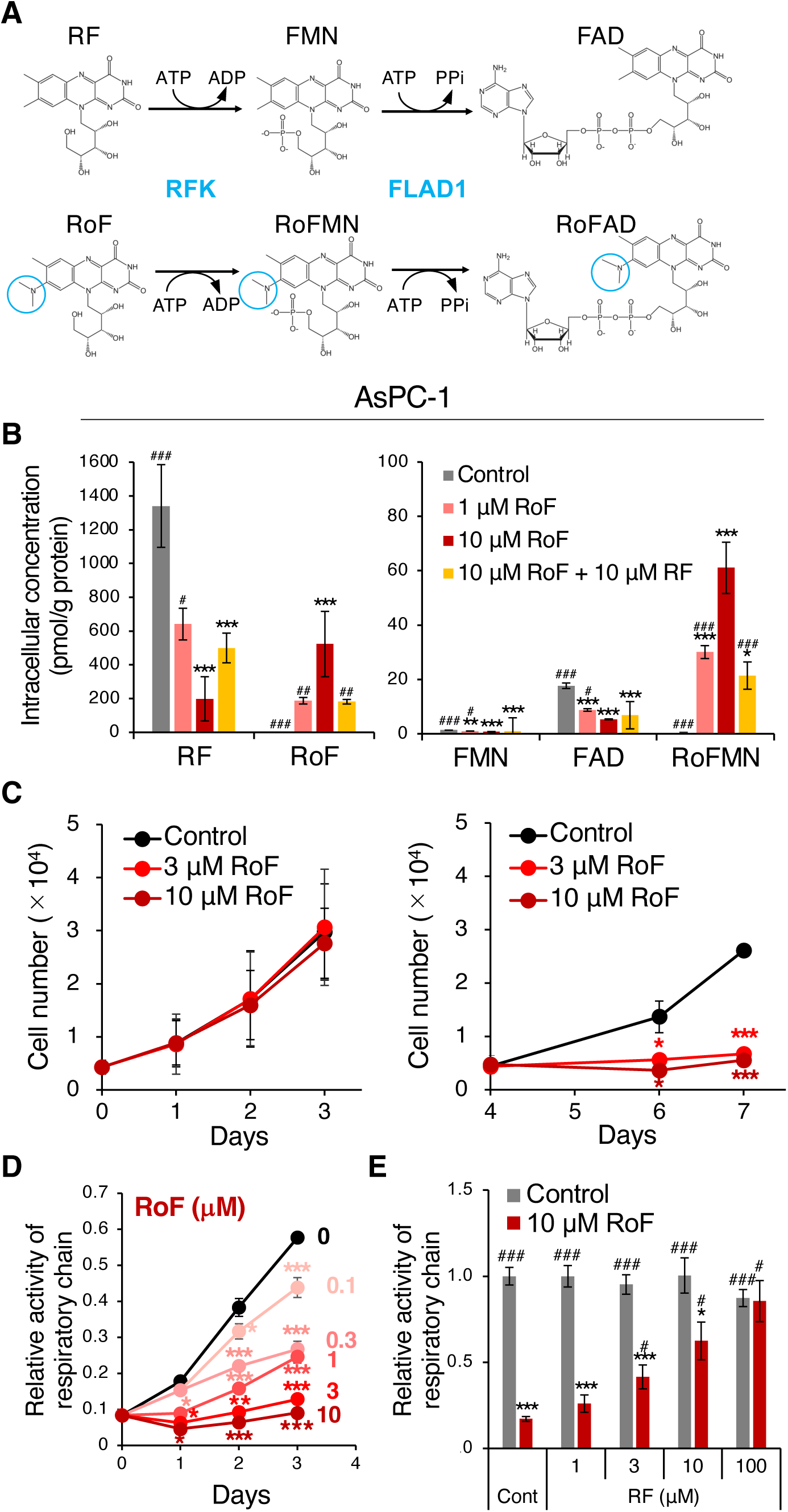
Roseoflavin treatment suppresses cell proliferation and mitochondrial respiratory chain activity in AsPC-1 cells by inhibiting RF metabolism. (A) RF and roseoflavin (RoF) are metabolized by RFK and FLAD1. RoF is an analog of RF, with the only structural difference being by the light blue circle. PPi, pyrophosphate. (B) RoF treatment decreased intracellular levels of RF and its metabolites by competing with RF. Metabolite levels were quantified 48 hours following treatment with vehicle, RoF, and/or RF. (C) Long-term treatment with RoF leads to a reduction in cell numbers. Viable cells were counted using trypan blue exclusion after treatment with vehicle or RoF. (D) RoF inhibited mitochondrial respiratory chain activity in a concentration-dependent manner. Activity was measured using an MTS assay kit. (E) RoF-mediated inhibition of mitochondrial respiratory chain activity was reversed by the addition of RF. Respiratory chain activity was assessed three days post-treatment. Error bars, SD in technical triplicate. *, *P* < 0.005; **, *P* < 0.0005; ***, *P* < 0.0001 in Dunnett’s test compared to control. ^#^, *P* < 0.05; ^##^, *P* < 0.005; ^##^, *P* < 0.0001 in Dunnett’s test compared to 10 μM RoF.

Therefore, we investigated the efficacy of RoF in inhibiting RF metabolism in PDAC cells. As a result, RoF treatment dose-dependently increased the levels of RoF and RoFMN in AsPC-1 cells, while simultaneously decreasing the levels of RF, FMN, and FAD (Fig. 2B). Furthermore, the addition of RF in the presence of RoF significantly suppressed the levels of RoF and RoFMN, and tended to restore the reduced levels of RF (Fig. 2B), suggesting that RoF inhibits RF metabolism by competing with RF in PDAC cells, consistent with findings reported in bacterial studies.

Moreover, we assessed the impact of RoF on the proliferation of AsPC-1 cells. Even though RoF treatment did not affect cell numbers within the first three days, reseeding these cells following RoF treatment significantly reduced their numbers (Fig. 2C). Additionally, RoF treatment suppressed respiratory chain activity in a dose-dependent manner (Fig. 2D), and this suppression was reversed by RF supplementation (Fig. 2E), and similar effects were observed in PANC-1 (Fig. EV2), MIA PaCa-2 (Fig. EV3A), and Capan-1 cells (Figs. EV3B and EV3C). Notably, RoF inhibited respiratory chain activity more potently than lumiflavin, another RF metabolism inhibitor (Figs. EV3B, EV3D, and EV3E). In respiratory chain, two flavoproteins, NADH:ubiquinone oxidoreductase core subunit V1 (NDUFV1) and Succinate dehydrogenase complex flavoprotein subunit A (SDHA), are involved in mitochondrial complexes I and II respectively. Therefore, these findings suggest that RoF treatment disrupts RF uptake and its metabolite conversion, leading to the inhibition of respiratory chain activity and cell proliferation.

### Combined inhibition of RF metabolism and MEK exhibits anti-tumor effects in a mouse xenograft model of human PDAC

Given that RoF inhibits RF metabolism and suppresses PDAC cell proliferation, we investigated whether RoF could similarly attenuate their proliferation in mice. To establish the appropriate injection dose of RoF for intraperitoneal administration, we determined the maximum tolerated dose < 100 mg/kg (Fig. EV4A). Furthermore, we conducted pharmacokinetic studies. Injection of mice with 10 mg/kg RoF resulted in a peak blood concentration of 8.48 μM (Fig. EV4B), which is comparable to the concentration that inhibited proliferation of cultured PDAC cells (Figs. 2B and EV2B). Based on our previous findings that therapeutic candidates identified in *4-hit* flies demonstrated enhanced effects when combined with trametinib (Tr) (Sekiya et al, 2023; Fukuda et al, 2024), we administered not only RoF but also Tr to potentially enhance anti-tumor effects. Monotherapy with either RoF or Tr did not induce tumor regression (Fig. 3A). However, the combination therapy significantly inhibited AsPC-1 xenograft growth compared to control group (Fig. 3A) and notably reduced tumor weight after four weeks of treatment compared to both control and Tr monotherapy (Fig. 3B). Importantly, the dosage of the combination treatment did not impact body weight of either non-transplanted or AsPC-1 tumor-bearing mice (Figs. EV4C and EV4D).

**Figure 3.**
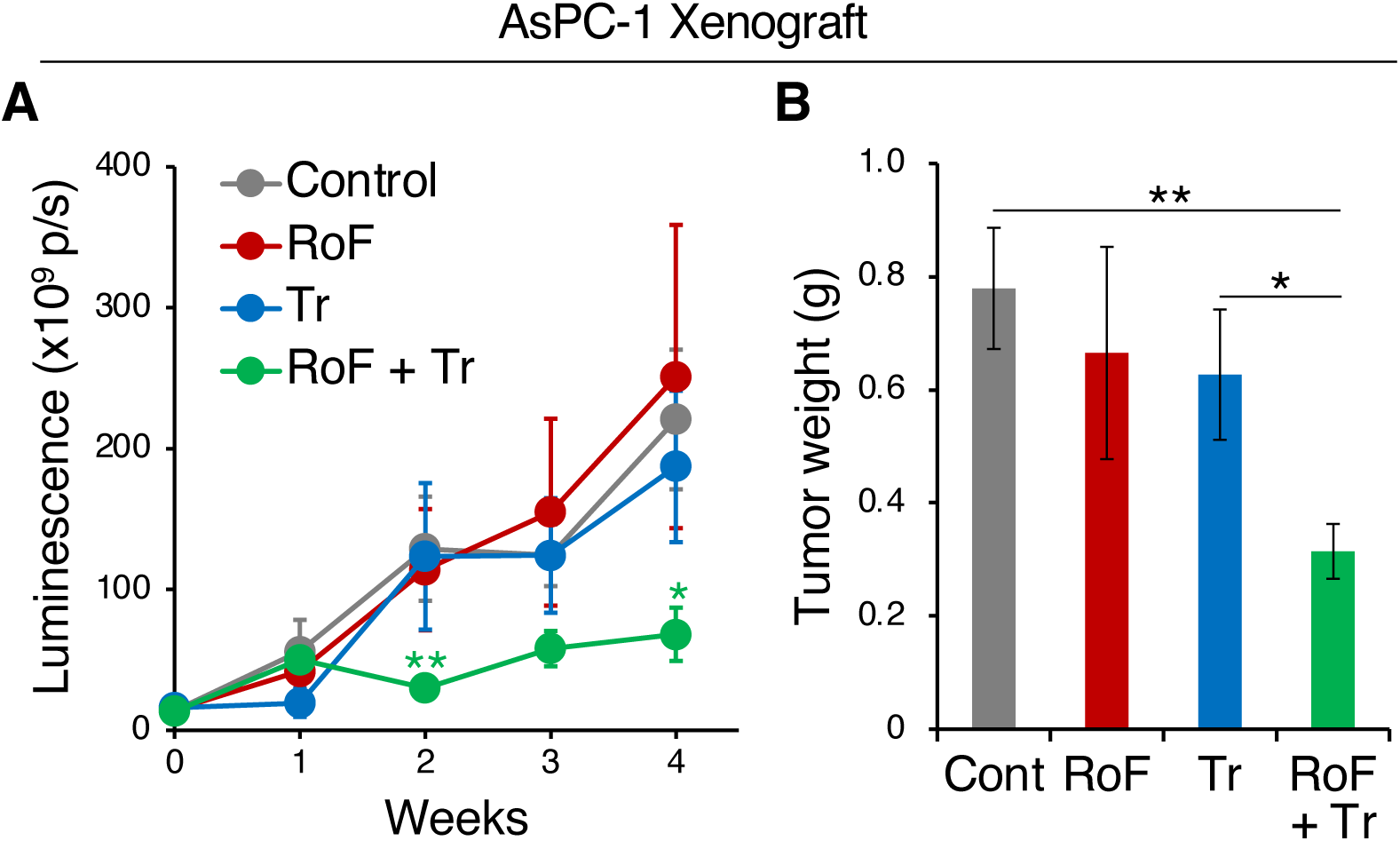
Combined inhibition of RF metabolism and MEK demonstrates a potent anti-tumor effect in orthotopic AsPC-1 xenografts in mice. (A) Treatment with a combination of RoF and trametinib (Tr) suppressed the expansion of AsPC-1 xenografts. Luciferase-expressing AsPC-1 xenografts were monitored by *in vivo* bioluminescent imaging. Mice were treated with vehicle (control), RoF alone (10 mg/kg, i.p.), Tr alone (0.2 mg/kg p.o.), and RoF + Tr combined (10 mg/kg RoF i.p. and 0.2 mg/kg Tr p.o.). Sample sizes at each time point were control (n = 7, 4, 6, 7, 7), RoF (n = 7, 5, 6, 7, 7), Tr (n = 6, 3, 6, 6, 6), and RoF + Tr (n = 6, 3, 6, 6, 6). Error bars, standard error. *, *P* < 0.05; **, *P* < 0.01 in the Mann-Whitney *U*-test compared to control. (B) Tumor weight, measured on day 28, showed a significant reduction following combination treatment compared to control or Tr treatment alone. Control and RoF (n = 7), Tr and RoF + Tr (n = 6). Error bars, SD. *, *P* < 0.05; **, *P* < 0.01 in the Mann-Whitney *U*-test.

Subsequently, four weeks after administering RoF and/or Tr, we analyzed the tumors using Western blotting to investigate the treatment effects. Contrary to expectations, phosphorylation levels of ERK were low or undetectable in control tumors, which precluded an assessment of Tr effects (Fig. EV5A). Moreover, treatments with RoF and/or Tr did not affect the levels of proliferation markers (phosphorylated histone H3 and PCNA, Fig. EV5A), an oxidative stress marker (γH2AX, Fig. EV5A), necroptosis markers (phosphorylated-RIP and -MLKL, Fig. EV5B), apoptosis markers (cleaved-PARP and Caspase-3, Fig. EV5B), or a pyroptosis marker (cleaved-GSDME, Fig. EV5B). This may be because the combined treatment with RoF and Tr does not affect PDAC cell proliferation, but rather induces cell death, leading to the rapid elimination of cells. These findings indicate that combined inhibition of RF metabolism and MEK represents a promising therapeutic strategy for PDAC.

### Combined inhibition of RF metabolism and MEK suppresses spheroid formation, potentially through inhibition of the TCA cycle

To evaluate whether the anti-tumor effects of RoF and Tr on directly target PDAC cells, AsPC-1 cells were cultured in three-dimensional (3D) media containing methylcellulose with added RoF and/or Tr. Spheroids were counted after three weeks following culture. Both RoF and Tr alone suppressed the formation of large spheroids compared to control (Fig. 4). Notably, their combination treatment significantly reduced the number of large, medium, and small spheroids (Fig. 4). Essentially, we obtained similar results for PANC-1 cells (Fig. EV6). These findings suggest that the anti-tumor effects of the combination treatment are, at least in part, directly at PDAC cells.

**Figure 4.**
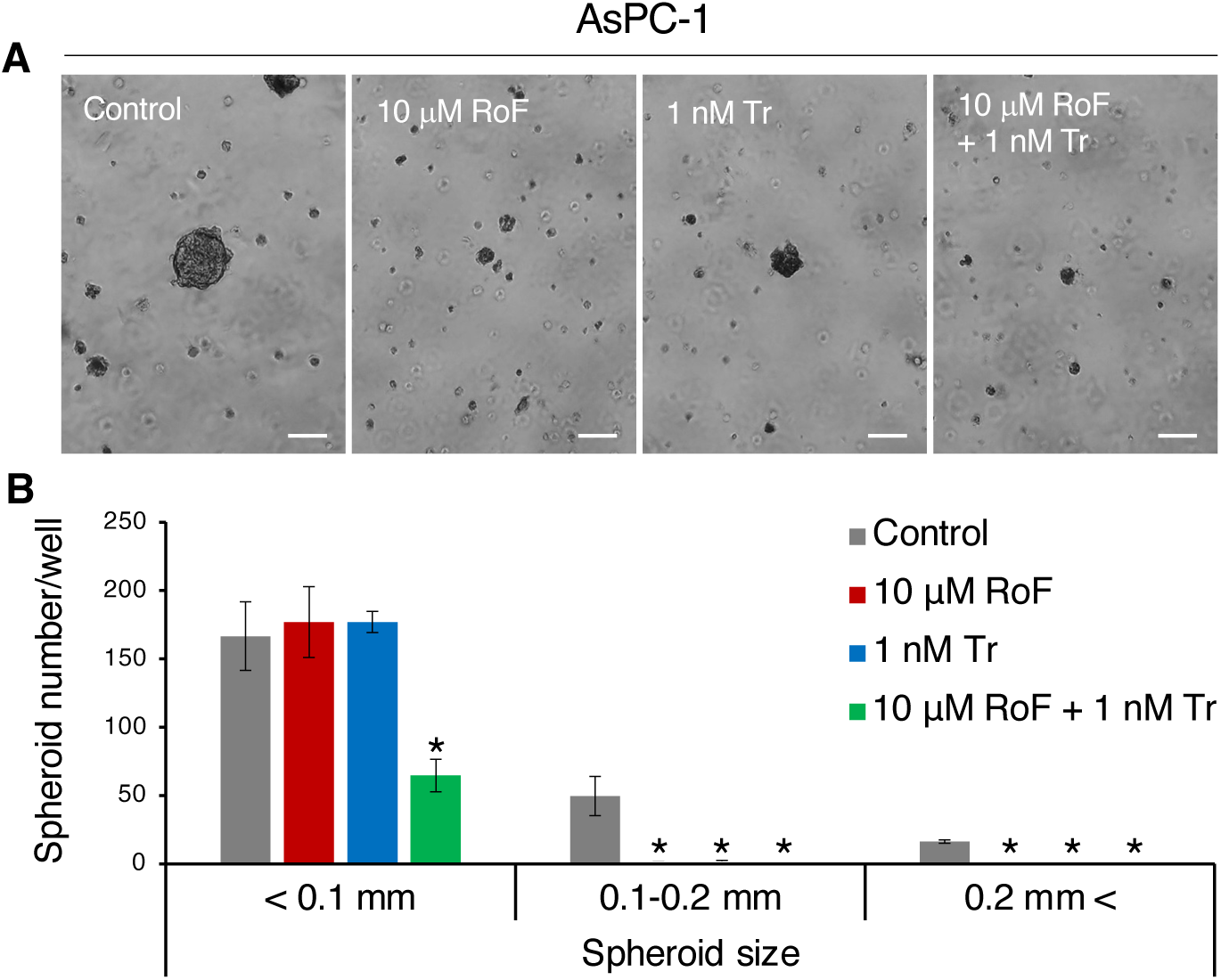
Inhibition of RF metabolism and MEK suppresses the formation of spheroids in AsPC-1 cells. (A) Three-dimensional cultures of AsPC-1 cells in methylcellulose media three weeks post-seeding. Scale bars, 0.1 mm. (B) Quantification of the number of spheroids by diameter size category following treatment. Error bars, SD in technical triplicate. *, *P* < 0.05 in Dunnett’s test compared to control.

Since RF metabolism regulates various metabolic processes, we conducted comprehensive (CE-TOFMS) and target (LC-MS/MS) metabolome analyses on 3D-cultured AsPC-1 and PANC-1 cells treated with RoF and/or Tr. Due to the low detection sensitivity of RF metabolites by CE-TOFMS, they were specifically analyzed using LC-MS/MS. These treatments resulted in significant changes in various metabolites compared to control (Tables 1, 2, EV1, and EV2). Similar to observations in 2D cultures, RoF or the combination treatment significantly decreased the levels of RF, FMN, and FAD in AsPC-1 cells (Fig. 5A). Additionally, 2-hydroxyglutarate (2-HG) levels increased under RoF or the combination treatment (Fig. 5A). PANC-1 cells also showed similar results (Fig. EV7A). Although mutation in *Isocitrate dehydrogenase [NADP] 1* (*IDH1*) or *IDH2* increases 2-HG levels (Dang et al, 2009; Ward et al, 2010), the Cancer Cell Line Encyclopedia (CCLE) database confirmed that the cell lines used in this study do not possess these mutations in *IDH1* or *IDH2*. Moreover, 2-HG is metabolized to α-ketoglutarate by L-2-hydroxyglutarate dehydrogenase (L2HGDH), which utilizes FAD as a coenzyme (Du et al, 2021). Therefore, the observed elevation in 2-HG levels, both with RoF alone and in combination with Tr, is potentially attributable to reduced intracellular FAD, inhibiting the activity of the flavoprotein L2HGDH.

**Figure 5.**
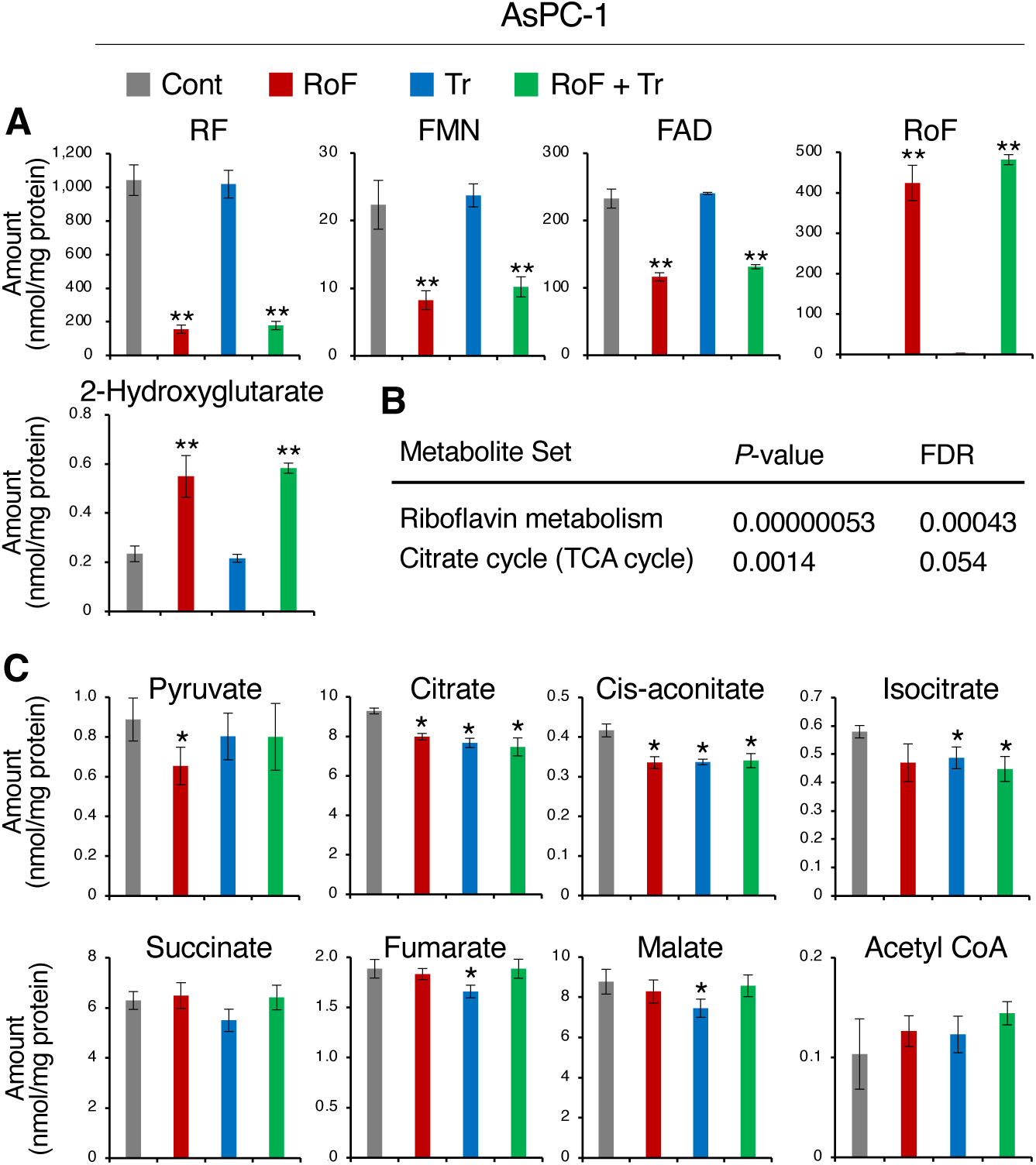
Combined inhibition of RF metabolism and MEK reduces metabolites in RF metabolism and the TCA cycle in AsPC-1 cells. (A) Treatment with RoF and/or Tr decreased intracellular levels of metabolites involved in RF. (B) Enrichment analysis indicated that combined treatment with RoF and Tr significantly reduced metabolites associated with RF metabolism and the TCA cycle. Metabolites showing a decrease of 1.2-fold or greater in cells treated with RoF and Tr compared to control cells were specifically analyzed using Metaboanalyst 6.0 (https://www.metaboanalyst.ca/). FDR, false discovery rate. (C) Treatment with RoF and/or Tr decreased intracellular levels of several TCA cycle-related metabolites. Error bars, SD in technical quadruplicate. **\*,** *P* < 0.05; **\*\*,** *P* < 0.01 in Dunnett’s test compared to control.

**Table 1.**
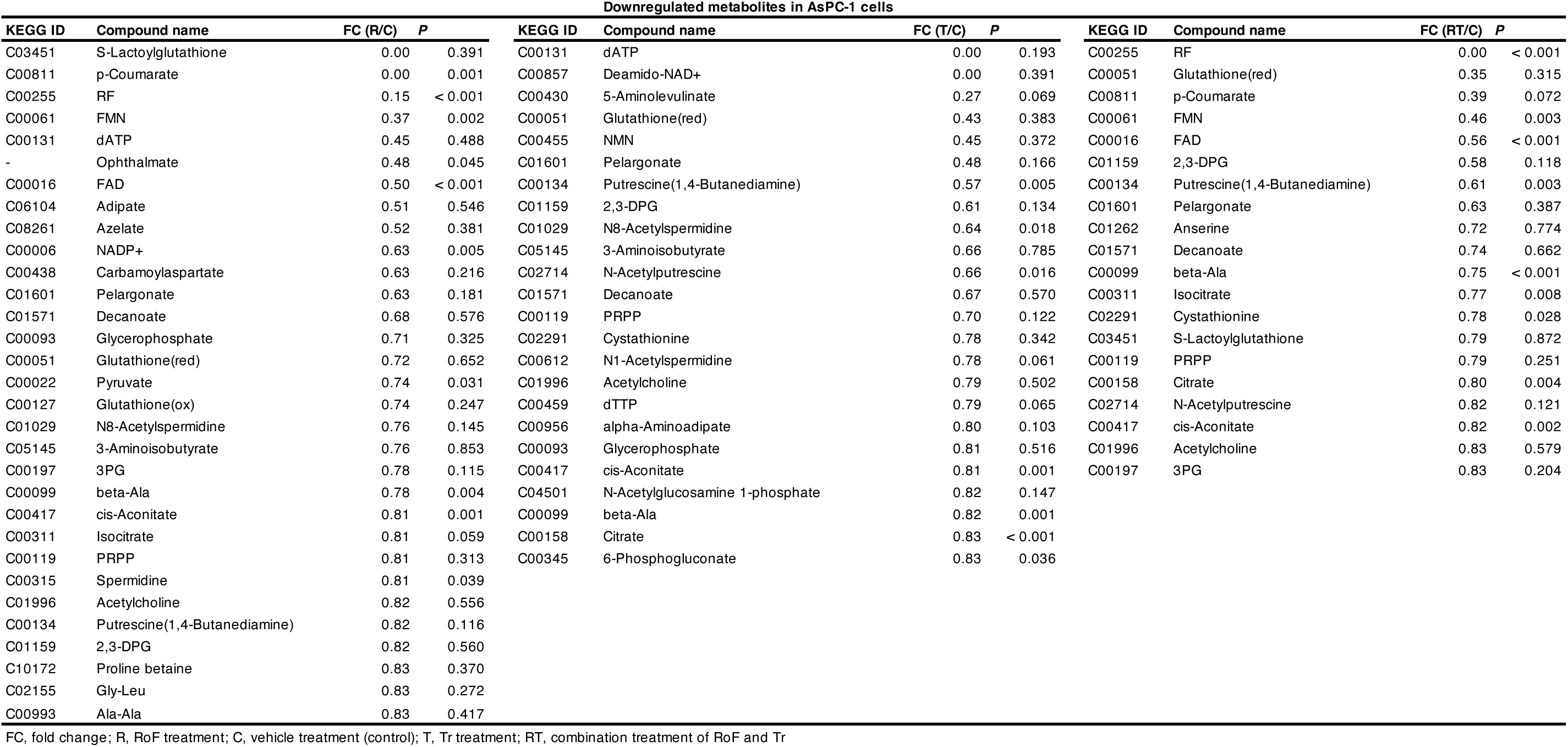
Metabolites reduced by 1.2-fold or more in AsPC-1 cells following treatment with RoF and/or Tr.

**Table 2.**
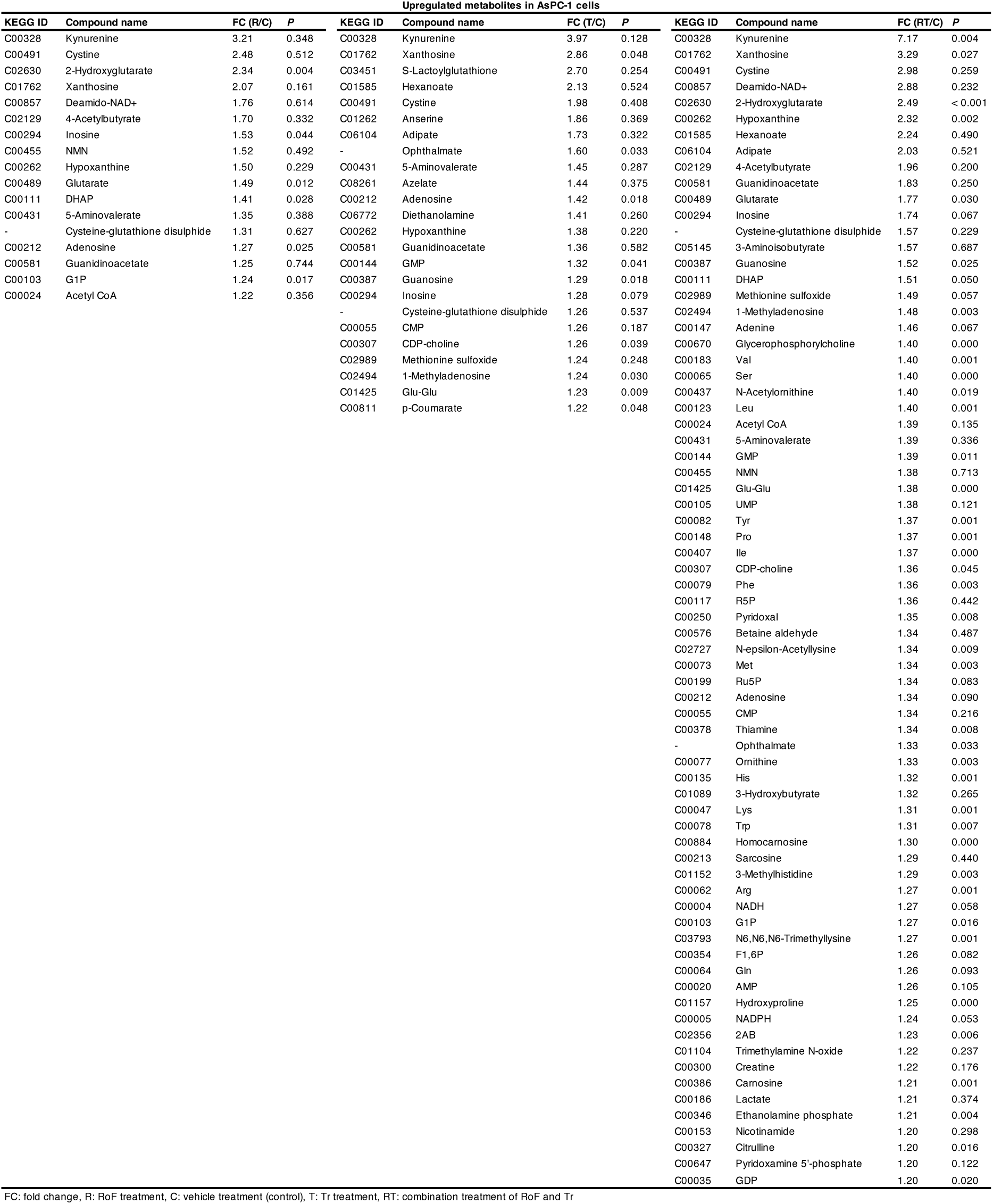
Metabolites elevated by 1.2-fold or more in AsPC-1 cells following treatment with RoF and/or Tr.

Furthermore, we observed distinct differences in the ratios of RF, FMN, and FAD between AsPC-1 and PANC-1 cells (Table EV3). In AsPC-1 cells, RF was the predominant metabolite among those measured, whereas FAD was the most abundant in PANC-1 cells (Figs. 6A and EV7A, Table EV3). Subsequent RNA-seq analysis of RF metabolism-related genes, conducted under identical culture conditions, revealed that *SLC52A2* had highest expression among RF transporter genes in both cell lines (Table EV3). Notably, *SLC52A2* expression in AsPC-1 cells was approximately threefold higher than in PANC-1 cells (Table EV3). Additionally, *RFK* expression in AsPC-1 was less than half that observed in PANC-1 cells (Table EV3). These results suggest that although AsPC-1 cells exhibit high RF levels, potentially due to elevated *SLC52A2* expression, the low expression of *RFK* may inhibit the conversion of RF to FMN and the subsequent synthesis of FAD compared to PANC-1 cells.

**Figure 6.**
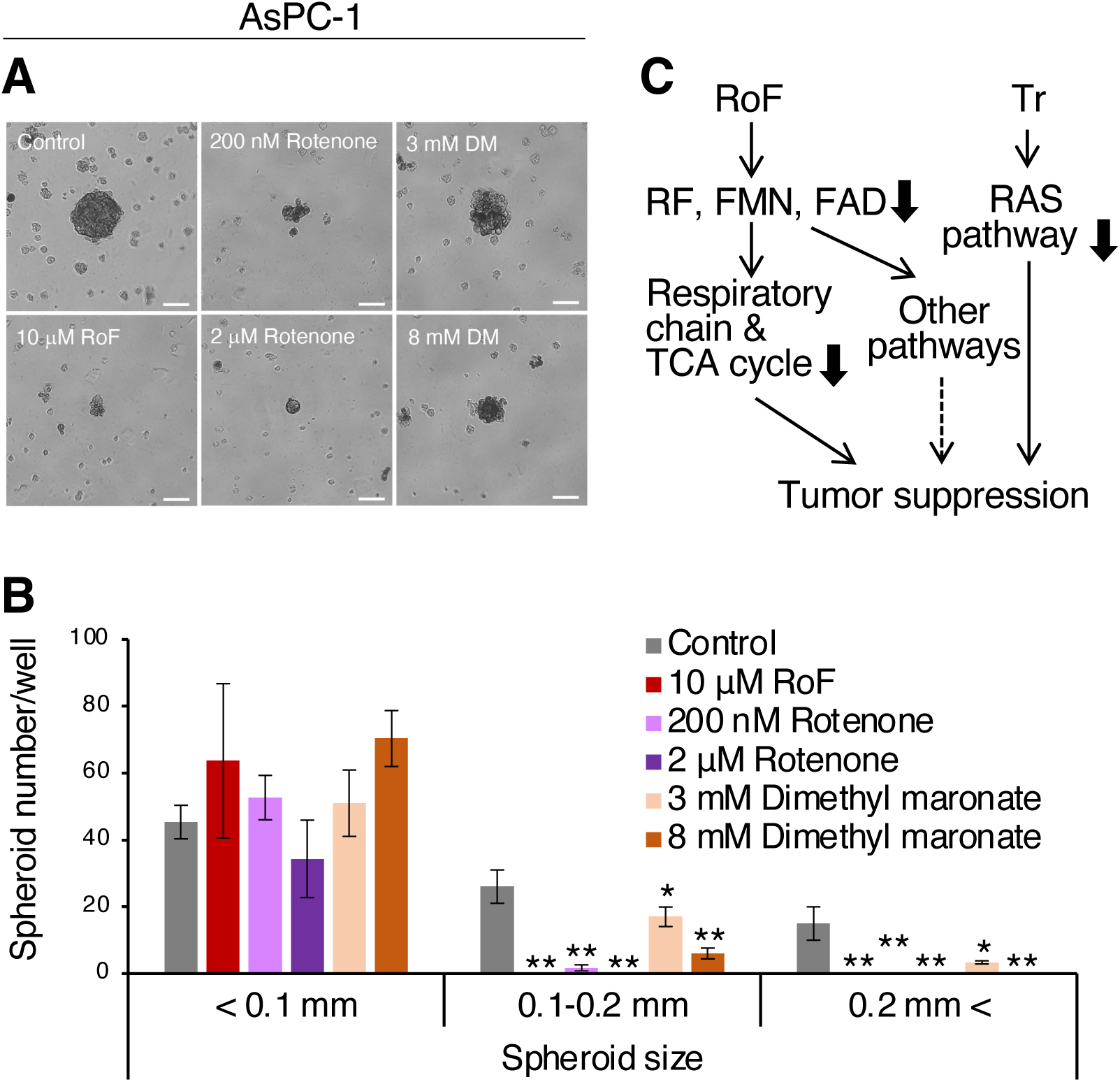
Inhibition of mitochondrial respiratory chain activity suppresses spheroid formation in AsPC-1 cells. (A) Three-dimensional cultures of AsPC-1 cells in methylcellulose media were treated with RoF, rotenone, or dimethyl malonate (DM) and analyzed three weeks post-seeding. Scale bars, 0.1 mm. (B) Quantification of spheroids by diameter size categories following treatment with inhibitors of mitochondrial respiratory chain activity. Error bars, SD in technical triplicate. *, *P* < 0.05 in Dunnett’s test compared to control. (C) A schematic representation of the mechanism of action of RoF and Tr.

To further investigate metabolic changes within the cells, we conducted enrichment analysis. This analysis revealed that metabolites decreased by more than 1.2-fold in RoF and Tr combination treatment compared to control were significantly enriched in pathways involved in RF metabolism and the tricarboxylic acid (TCA) cycle in AsPC-1 cells (Fig. 5B). Examination of TCA cycle metabolites in these cells showed significant reductions in citrate, cis-aconitate, and isocitrate levels following the combination treatment (Fig. 5C). Conversely, enrichment analysis in PANC-1 cells indicated that the combination treatment reduced metabolites related to RF metabolism and several amino acid pathways (Fig. EV7B). Specifically, the levels of serine, tyrosine, valine, leucine, histidine, phenylalanine, methionine, and threonine were decreased by RoF and/or Tr treatment in PANC-1 cells (Fig. EV7C). Interestingly, these amino acid levels increased under combination treatment in AsPC-1 cells (Fig. EV8A), suggesting that the impact of the combination treatment on amino acid metabolism varies between cell types.

Additionally, levels of pyruvate and succinate, which are associated with the TCA cycle, were decreased by the combination treatment (Fig. EV8B). The TCA cycle is associated with the flavoprotein SDHA (Porporato et al, 2018). Despite these metabolic reductions, the expression of genes related to the TCA cycle was not decreased by RoF and/or Tr treatment (Figs. EV9A and 9B), suggesting that the decline in TCA cycle metabolites may be due to decreased enzyme activity of SDHA. These results are consistent with our findings of reduced respiratory chain activity following RoF treatment, which correspondingly diminished TCA cycle activity in both AsPC-1 and PANC-1 cells.

Furthermore, the amount of aspartate decreased in both AsPC-1 and PANC-1 cells following RoF treatment (Figs. EV10A and EV11A). Previous studies have shown that cell proliferation by inhibition of respiratory chain, can be restored by supplementing the medium with pyruvate or aspartate (Birsoy et al, 2015; Sullivan et al, 2015). Considering that RoF treatment reduces respiratory chain activity (Figs. 2C and EV2C), we hypothesized that inhibitory effect of RoF treatment on cell proliferation may be due to a decrease in pyruvate or aspartate. However, the addition of pyruvate or aspartate failed to reverse the RoF-induced suppression of spheroid formation and respiratory chain activity (Figs. EV10B, 10C, 10D, EV11B, 11C, and 11D), despite the abundant expression of pyruvate and aspartate transporters in both AsPC-1 and PANC-1 cells (Figs. EV10E and EV11E). In other words, the supplementation of pyruvate or aspartate was not sufficient to counteract the reduction in spheroid formation ability caused by RoF treatment. These findings suggest that the inhibition of spheroid formation by RoF is mediated by a mechanism independent of pyruvate or aspartate inhibition.

Finally, to determine whether the anti-tumor effects of RoF is due to inhibition of the respiratory chain, we added inhibitors of mitochondrial complex I or II, rotenone or dimethyl malonate, to 3D-cultured AsPC-1 and PANC-1 cells. As a result, spheroid formation was inhibited similarly to RoF treatment (Figs. 6A, 6B, EV12A, and EV12B). Furthermore, we did not observe inhibition of RAS pathway in Tr-treated xenografts. However, JFCR LinCAGE analysis using RNA-Seq revealed that the gene signatures of Tr-treated spheroids closely resembled those of cells treated with MEK (U-0126) and BRAF (dabrafenib) inhibitors (Table EV4). These results suggest that RoF inhibits the respiratory chain and other flavoprotein-involved pathways by reducing intracellular RF and its metabolites (Fig. 6C). Additionally, Tr enhances its anti-tumor effects by inhibiting RAS pathway (Fig. 7C).

## Discussion

In this study, genetic screening utilizing in *4-hit* flies identified RF metabolism as a novel therapeutic target for PDAC at the whole-body level. Moreover, combining the RF metabolism inhibitor RoF with the MEK inhibitor Tr effectively suppressed the growth of human PDAC xenografts and 3D-cultured spheroids. Mechanistically, we found that RoF treatment not only reduced the levels of RF, FMN, FAD, and several TCA cycle-related metabolites, while also reducing respiratory chain activity in both AsPC-1 and PANC-1 cells. Interestingly, the responses of amino acid metabolites to RoF treatment varied between these cells, suggesting that different PDAC cell subtypes may exhibit unique amino acid metabolic profiles. In a metabolic study of PDAC cells, a study categorized PDAC cell lines into three metabolic subtypes—slow proliferating, glycolytic, and lipogenic—based on metabolite profiles (Daemen et al, 2015), AsPC-1 cells were classified as the slow proliferating subtype, while PANC-1 cells were categorized as the lipogenic subtype. Our findings align with this classification; amino acid levels were higher in PANC-1 cells than in AsPC-1 cells, consistent with our metabolomic results. In PANC-1 cells, respiratory chain and TCA cycle activities were reduced by RoF treatment, potentially prompting the use of amino acids to compensate for the energy deficit. Furthermore, MIA PaCa-2 cells, classified as glycolytic, exhibit a metabolism that relies more on glycolysis than on the respiratory chain (Daemen et al, 2015). Consistent with this, our data indicated that the inhibitory effect of RoF on respiratory chain inhibitory effect of RoF in MIA PaCa-2 cells was less pronounced compared to other PDAC cell lines. These results suggest that RoF treatment may be particularly effective in cells with high respiratory chain activity, regardless of their specific gene mutation profile. Although RF metabolism was initially identified through screening in *4-hit* flies, the efficacy of the RoF and Tr combination in PANC-1 cells, which harbor *3-hit* alterations in *KRAS*, *TP53*, and *CDKN2A* (CCLE database and Sekiya et al, 2023).

Research into the regulatory mechanisms of RF metabolism has advanced significantly in bacteria. In bacteria, when excessive FMN is produced within the cells, it binds to the FMN riboswitch, suppressing the expression of genes such as RFT and thereby inhibiting the uptake of RF (Mansjo and Johansson, 2011). However, riboswitch has not been discovered in mammals, and the regulatory mechanisms of RF metabolism remain unclear. Nevertheless, by comparing the measurement of RF metabolites and RNA-Seq results in AsPC-1 and PANC-1 cells, we have identified a potential regulation of RF metabolism through the expression of RF metabolism-related genes. Furthermore, we have demonstrated that the expression of genes related to RF metabolism was upregulated in PDAC compared to non-tumorous pancreatic tissue, suggesting that RF metabolism may be activated in PDAC. Therefore, the inhibition of RF metabolism in PDAC suggests a potential to preferentially hinder its growth.

Furthermore, we employed RoF to inhibit RF metabolism. In bacteria, RoF is metabolized into RoFMN and RoFAD by RFK and FLAD1, respectively (Fig. 2A). RoFMN binds to the FMN riboswitch, effectively suppressing the levels of RF metabolites, even in the presence of insufficient FMN within bacteria (Mansj and Johansson, 2011). Thus, RoFMN can serve as an alternative to inhibit RF metabolism. Although this regulatory mechanism is well-documented in bacteria, the effects of RoF in mammalian cells were previously unclear. We have now elucidated, for the first time, the impact of RoF on PDAC cells, revealing that RoF was metabolized RoFMN and RoF treatment reduced the levels of RF, FMN, and FAD. Previously, lumiflavin (LF) is utilized to inhibit RF metabolism in mammalian cells (Yang et al, 2019). However, our studies demonstrate that RoF exerted a more potent inhibitory effect on activity of the respiratory chain, which operates downstream of RF metabolism, compared to LF. LF is deficient in the ribityl chain required for phosphorylation by RFK (Fig. EV3E). Therefore, the significant inhibitory impact of RoF on RF metabolism may originate from its competitive inhibition of RFK and FLAD1, which metabolize RF and FMN, respectively.

Notably, RoFMN does not function as a cofactor for the flavoprotein D-amino acid oxidase (DAAO) in a cell-free assay (Grill et al., 2008), suggesting that RoFMN cannot activate flavoenzymes. This evidence suggests that RoFMN produced in mammalian cells competitively inhibits the activity of FMN-binding flavoproteins.

RF metabolism has been reported to influence various metabolic pathways and antioxidant activities (Joosten and Berkel, 2007). Enzymes that utilize FMN and FAD as cofactors are known as flavoproteins, and 111 types have been identified to date (Wegrzyn et al, 2019). Among these, we hypothesized that the anti-tumor effects of RoF may be due to the inhibition of SDHA and HDUFV1, which act in the respiratory chain. Because significant inhibition of respiratory chain activity was consistently observed across various PDAC cell lines following RoF (Figs. 2C, EV2C, EV3A, and EV3B). Indeed, targeting the respiratory chain has been proposed as a therapeutic strategy in multiple types of cancers, including PDAC (Dong et al, 2020; Porporato et al, 2018). Specifically, in PDAC, inhibitors of mitochondrial complex I, such as phenformin and metformin, have demonstrated anti-tumor effects, which are enhanced when combined with the PDAC drug gemcitabine (Masoud et al, 2020; Suzuki et al, 2019). However, there have been no reports on the co-inhibition of the respiratory chain and MEK as a cancer therapeutic approach. In this study, we demonstrated that the combined inhibition of RF metabolism and MEK produced significantly greater anti-tumor effects than either inhibition alone. In contrast, previous research on prostate and ovarian cancers that targeted RF metabolism as a therapeutic strategy combined its inhibition with cisplatin (Hirano et al, 2011; Yang et al, 2019). Therefore, our approach of simultaneously inhibiting RF metabolism and MEK represents a novel strategy in cancer research. Given that over 90% of PDAC cases harbor *KRAS* mutations (Morris et al, 2010), MEK inhibitors may be particularly effective for PDAC, potentially offering a more specific and less toxic alternative to the combination of DNA synthesis inhibitors like gemcitabine and cisplatin. While we focused on the respiratory chain as a downstream of RF metabolism, it is possible that inhibition of other flavoproteins may also contribute to the observed anti-tumor effects. Further research is required to elucidate these potential interactions.

In summary, our study provides compelling evidence that dual inhibition of RF metabolism and MEK signaling represents a promising therapeutic approach for PDAC. By targeting these two critical pathways, we introduce a novel paradigm that combines the inhibition of protein kinases and vitamin kinases, aiming to reduce the resilience of PDAC cells more effectively. Elucidating the intricate role of RF metabolism in regulating oncogenic signals will shed light on the broader implications of how essential nutrients, commonly found in the diet, contribute to PDAC. This could lead to a deeper understanding of the molecular mechanisms driving PDAC progression and provide valuable insights into new therapeutic interventions that exploit these vulnerabilities.

## Methods

### Reagents

See Table 3.

**Table 3.** Reagents.

|  |
| --- |
| 2.5g/l-Trypsin/1mmol/l-EDTA Solution, Nacalai Tesque, Kyoto, Japan, 35554-64 |
| ReverTra Ace qPCR RT Master Mix, Toyobo, Japan, QPS-201 |
| Thunderbird SYBR qPCR Mix, Toyobo, Japan, QPS-201 |
| Riboflavin, Tokyo Chemical Industry, Tokyo, Japan, R0020 |
| Riboflavin, Fujifilm Wako Pure Chemical Corporation, Osaka, Japan, 180-00171 |
| Flavin mononucleotide, Tokyo Chemical Industry, Tokyo, Japan, R0023 |
| Flavin adenine dinucleotide, Tokyo Chemical Industry, Tokyo, Japan, F0014 |
| Roseoflavin mononucleotide, homemade, refer to Pedrolli et al, 2011 |
| 0.1% formic acid in water, Fisher Chemical, USA, LS118-1 |
| 0.1% formic acid in acetonitrile, Fisher Chemical, USA, LS120-1 |
| Lumiflavin, Toronto research chemicals, North York, Canada, L473900 |
| D-Luciferin Potassium Salt, Fujifilm Wako Pure Chemical Corporation, Osaka, Japan, 126-05116 |
| SDS-PAGE sample loading buffer [6x] , G-Biosciences, St. Louis, USA, 786-701 |
| 2-Mercaptoethanol, Bio-rad, California, USA, 1610710 |
| 30% Acrylamide/Bis Solution, Bio-rad, California, USA, 1610158 |
| Ammonium peroxodisulfate, Fujifilm Wako Pure Chemical Corporation, Osaka, Japan, 012-08023 |
| Tris (hydroxymethyl) aminomethane, Nacalai Tesque, Kyoto, Japan, 35406-75 |
| Sodium Dodecyl Sulfate, Fujifilm Wako Pure Chemical Corporation, Osaka, Japan, 192-13981 |
| N,N,N',N'-Tetramethylethylenediamine, Fujifilm Wako Pure Chemical Corporation, Osaka, Japan, 205-06313 |
| Polyoxyethylene(20) sorbitan monolaurate, Fujifilm Wako Pure Chemical Corporation, Osaka, Japan, 166-2111 |
| 10xTBS, Bio-rad, California, USA, 1706435 |
| Anti-Phospho-p44/42 MAPK (Erk1/2) (Thr202/Tyr204) (D13.14.4E) XP® Rabbit mAb, Cell Signaling Technology |
| Anti-p44/42 MAPK (Erk1/2) (137F5) Rabbit mAb, Cell Signaling Technology, Boston, US, 4370, AB_2315112 |
| Anti-Phospho-Histone H3 (Ser10) Antibody, Cell Signaling Technology, Boston, US, 9701, AB_331535 |
| Anti-Histone H3 (D1H2) XP® Rabbit mAb, Cell Signaling Technology, Boston, US, 4499, AB_10544537 |
| Anti-PCNA (D3H8P) XP® Rabbit mAb, Cell Signaling Technology, Boston, US, 13110, AB_2636979 |
| Anti-Phospho-Histone H2A.X (Ser139) (20E3) Rabbit mAb, Cell Signaling Technology, Boston, US, 9718, AB_2 |
| Anti-α-Tubulin (11H10) Rabbit mAb, Cell Signaling Technology, Boston, US, 2125, AB_2619646 |
| Anti-β-Actin Antibody, Cell Signaling Technology, Boston, US, 4967, AB_330288 |
| Anti-Phospho-RIP3 (Ser227) (D6W2T) Rabbit mAb, Cell Signaling Technology, Boston, US, 93654, AB_280020 |
| Anti-Phospho-MLKL (Ser358) (D6H3V) Rabbit mAb, Cell Signaling Technology, Boston, US, 91689, AB_289504 |
| Anti-PARP (46D11) Rabbit mAb, Cell Signaling Technology, Boston, US, 9532, AB_659884 |
| Anti-Gasdermin E (E2X7E) Rabbit mAb, Cell Signaling Technology, Boston, US, 19453 |
| Anti-Caspase-3 Antibody, Cell Signaling Technology, Boston, US, 9662, AB_331439 |
| Anti-rabbit IgG, HRP-linked Antibody, Cell Signaling Technology, Boston, US, 7074, AB_2099233 |
| Immobilon Forte Western HRP substrate, MERCK, WBLUF0500 |
| Rotenone, MedChemExpress, HY-B1756 |
| Dimethyl malonate, MedChemExpress, HY-1787 |

### *Drosophila* genetic testing

To model the four gene alterations observed in PDAC patients, we have developed *4-hit* fly (Sekiya et al, 2023). This model involves the induced overexpression of *Drosophila Ras*^G12D^ (d*Ras*^G12D^) and *Cyclin E* (d*CycE*) and knockdown sequences of short hairpin RNA (shRNA) for *p53* and *Med*. Fly stocks harboring specific gene mutations or RNA interference (RNAi) were acquired from several resource centers, such as Bloomington *Drosophila* Stock Center (BDSC, Bloomington IN, USA), the National Institute of Genetics (NIG, Mishima, Japan), and Vienna Drosophila Resource Center (VDRC, Vienna, Austria). Fly stocks initially balanced with balancer chromosomes lacking *Tubby* (*Tb*) gene were rebalanced with *FM7c*-*Tb*-*RFP*, *CyO*-*Tb*-*RFP* (Pina and Pignoni, 2012), or *TM6B* balancer carrying *Tb* as a discernible marker. To knockout or knockdown a gene in the transformed cells, *Ser*>*Ras*^G12D^,*UAS*-*p53*^shRNA^,*UAS*- *dCycE*,*UAS*-*Med*^shRNA^/*SM5*_tub-GAL80_-*TM6B* (*Ser*>*4-hit*/*SM5*_tub-GAL80_-*TM6B*) flies were crossed with the balanced stocks containing either a specific gene mutation or a *UAS*-RNAi construct (see Table EV5). Control crosses for these experiments were performed by mating *4-hit* flies with nontransgenic flies (*w*^−^, BDSC, #3605). The resulting offspring were cultured until adulthood for a period of 11 days at 27°C, and their viability was assessed by calculating the ratio of the number of empty pupae to the total number of non-*Tb* pupae.

### Cell lines and cell culture

Human PDAC cell lines, PANC-1 and MIA PaCa-2, were obtained from RIKEN BRC CELL BANK (Tsukuba, Japan). Authenticated Capan-1 and AsPC-1 were purchased from ATCC. All cell lines were cultured in appropriate media (Table EV6) supplemented with fetal bovine serum (Nichirei Bioscience #175012, Tokyo, Japan) and 1% penicillin/streptomycin (Nacalai Tesque #26253-84, Kyoto, Japan) at 37°C in a humidified atmosphere with 5% CO_2_. All cell lines were utilized within 2 months post-thawing.

### Measuring riboflavin and its metabolites

Subconfluent PDAC cells were cultured in 10-cm dishes for 48 hours in media containing either vehicle (dimethyl sulfoxide, DMSO; Sigma-Aldrich #D2650, Madrid, Spain), 10 µM roseoflavin (RoF, Toronto Research Chemicals #R685000, Toronto, Canada), and/or 1 nM trametinib (Tr, MedChemExpress #HY-10999, Shanghai, China). Following incubation, cells were washed with PBS and collected using a scraper into 10 mL PBS. The cell suspension was centrifuged at 180 × *g* for 5 min at 4°C. The supernatant was aspirated, and the cell pellet was resuspended in 1 mL PBS and transferred to Eppendorf tubes. These tubes were further centrifuged at 860 × *g* for 5 min at 4°C, and the supernatant was discarded. This centrifugation step was repeated to maximize liquid removal, and the cell pellets were subsequently stored at -80°C.

For cell lysis, the pellet was resuspended in 260 µL of Milli-Q water and sonicated in an ice-water bath for 30 min. Following sonication, the suspension was centrifuged at 13,800 × *g* for 5 min at 4°C, and the supernatant was collected into a new tube. Protein concentrations were determined using Quick Start Bradford 1× dye (Bio-Rad #5000205, California, USA), and sample concentrations were normalized. Each cell lysate (250 µL) was mixed with 1 mL of cold methanol (Fujifilm Wako Pure Chemical #137-01823, Osaka, Japan) and 1 ng of isotopically labeled riboflavin (RF ^13^C_4_,^15^N_2_ from Toronto Research Chemicals #V676023), and incubated at -80°C for over 30 min. After incubation, samples were centrifuged at 18,800 × *g* for 20 min at 4°C. The supernatant was transferred to new tubes and evaporated under vacuum. The dry samples were stored at -20°C until mass spectrometry analysis.

We measured RF and its metabolites in the cells by liquid chromatography mass spectrometry (LC-MS) analysis. The dried extracts were redissolved in 100 μL of water. After sonication, the suspension was centrifuged at 13,800 × *g* for 5 min at 4°C, and the supernatant was filtered with a 0.45 μm syringe filtration disk to the vials for injection in the following LC-MS system. The resulting samples (5 μL) were analyzed using LC (LC-20AD UFLC system, Shimadzu, Kyoto, Japan) coupled to a triple quadrupole mass spectrometer (LCMS-8040, Shimadzu). A chromatographic separation was performed on an L-column 3 C18 (2.0 mm i.d. x 150 mm, 3 μm particle size, metal-free, CERI, Tokyo, Japan) at 40°C. The mobile phases consisted of a linear gradient of water (A) and acetonitrile (B) containing both 0.1% (v/v) formic acid: A/B = 98:2 (v/v) at 0 min; 48:52 (v/v) at 10 min. The flow-rate was 200 μL/min. The effluent from the column was measured by MS using electrospray ionization (ESI). ESI parameters were as follows: DL temperature 180°C; heat-block temperature 400°C; drying gas flow 15 L/min; and ESI voltage 4.5 kV. The mass spectrometer was operated in a multiple reaction monitoring (MRM) positive ionization mode. For analysis, the monitored transitions for RF, FMN, FAD, RoF, RoFMN and isotopically labeled RF were *m/z* 377.1 → 243.0 (elution time: 6.6 min), *m/z* 457.1 → 439.1 (elution time: 6.2 min), *m/z* 786.2 → 348.1 (elution time: 5.8 min), *m/z* 406.1 → 272.0 (elution time: 6.7 min), *m/z* 486.1 → 468.1 (elution time: 6.3 min), and *m/z* 380.1 → 246.0 (elution time: 6.6 min), respectively.

### Cell proliferation assay

We defined the day of compound treatment on PDAC cells as Day 0 and subsequently performed temporal cell counting. Specifically, PDAC cells were seeded at densities of 3,000 or 6,000 cells per well in 48-well plates for cell counting on Days 0-3 and 25,000 or 50,000 cells per well in 6-well plates for counting on Days 4-7. The following day, compounds were dissolved in 10% of the medium volume and added to each well, achieving a final DMSO concentration of 0.1%. Cell numbers for Days 0-3 were determined in triplicate by counting trypan blue (Fujifilm Wako Pure Chemical Corporation #207-17081)-positive cells. On Day 3, cells from the 6-well plates were reseeded at 3,000 or 6,000 cells per well into 48-well plates, treated with the compound-containing medium, and cell counts for Days 4-7 were conducted similarly.

### MTS assay

Between 1,000 and 3,000 PDAC cells in 100 μL culture medium were incubated in triplicate in 96-well plates for 24 hours. Subsequently, 11 μL of media containing compounds dissolved in DMSO were added to each well, achieving a final DMSO concentration of 0.1%. Post-incubation, mitochondrial respiratory chain activity was measured daily using CellTiter 96 AQueous One Solution Reagent (Promega #G3580, Madison, USA) and a microplate reader (iMark, Bio-Rad), following the manufacturer’s instructions. The activity was calculated relative to vehicle control.

### Mouse MTD assay

All mice were housed under specific pathogen-free conditions on a 12:12 hour light:dark cycle. To determine MTD for RoF, female BALB/c-*nu*/*nu* mice (6- or 12-week-olds; CLEA, Tokyo, Japan) were administered with increasing intraperitoneal (i.p.) doses of RoF, starting at 2 mg/kg/day. The mice were observed for one day for signs of clinical distress, such as weight loss, discharges, and morbidity. The dose was gradually escalated up to 100 mg/kg/day.

### Mouse pharmacokinetic assay

Pharmacokinetic assays for RoF were conducted by Medicilon (Shanghai, China). Six-week-old male ICR mice (Sino-British SIPPR/BK Lab Animal Ltd, Shanghai, China) were administered 10 mg/kg of RoF i.p., and the plasma concentrations of RoF were measured at 0.083, 0.25, 0.5, 1, 2, 4, 8, and 24 hours after dosing.

### Mouse xenograft assay

All surgical procedures were conducted under anesthesia, induced by subcutaneous administration of medetomidine (15 mg/kg, Kyoritsu Seiyaku, Tokyo, Japan), midazolam (Dormicum Injection, 200 mg/kg, Astellas Pharma, Tsukuba, Japan), and butorphanol tartrate (250 mg/kg, Meiji Seika Pharma, Tokyo, Japan). Post-procedure, anesthesia was reversed using atipamezole (150 mg/kg, Kyoritsu Seiyaku).

AsPC-1 cells stably expressing luciferase (hereafter referred to as AsPC-1-Luc) were generated as previously described (Sekiya et al, 2023). A total of 1,000,000 AsPC-1-Luc cells were injected into the exposed pancreatic tails of anesthetized female BALB/c-*n*u/*nu* mice (10- week-olds). Following the injection, the mice were randomly allocated to one of four groups based on similar average tumor size, determined using bioluminescence measured with IVIS Spectrum Imaging System (Caliper Life Science, Mountain View, USA). Treatments were administered five times per week for four weeks, with the mice receiving either vehicle (5% DMSO in saline, orally [p.o.]) or trametinib (Tr, 0.2 mg/kg p.o.), combination with vehicle (25% Cremophor EL [Nacalai Tesque #09727-14] and 25% ethanol (Fujifilm Wako Pure Chemical, #057-00451) in saline i.p.) or RoF (10 mg/kg i.p.). Weekly bioluminescence signals were captured using the IVIS system and quantified with Living Image Ver. 4.2 (Caliper, Hopkinton, USA).

### Western blotting

Excised xenograft tumors were cut into small pieces, immediately frozen in liquid nitrogen, and stored at -150°C. To extract proteins, deep-frozen samples were completely homogenized using BioMasher II (Nippi, Tokyo, Japan) in Cell Lysis Buffer (Cell Signaling Technology #9803, Danvers, USA) containing phenylmethylsulfonyl fluoride (Fujifilm Wako Pure Chemical Corporation #195381). Protein concentrations were determined using Quick Start Bradford 1× dye (Bio-Rad #5000205). Subsequently, proteins were separated by 10-15% SDS-PAGE and transferred to polyvinylidene difluoride membranes (Millipore #ISEQ07850, Billerica, USA), followed by blocking with Blocking One (Nacalai Tesque #03953-95) or Blocking One P (Nacalai Tesque #05999-84) for 30 or 20 min, respectively. The membranes were incubated with primary antibodies overnight at 4°C and then probed with secondary antibodies: horse anti-mouse IgG (Cell Signaling Technology #7076) or goat anti-rabbit IgG (Cell Signaling Technology #7074). Immunoreactivity was analyzed using ChemiDoc XRS+ and Image Lab software (Bio-Rad).

### Spheroid formation assay

To prepare methylcellulose stock solution, we followed the procedure described previously (Ware et al, 2016). Initially, autoclaved methyl cellulose 4,000 (Fujifilm Wako Pure Chemical #136-02155) was mixed with half the volume of the culture medium and stirred for 20 min while being heated at 60°C. Subsequently, the remaining half of the medium was added, and the mixture was stirred overnight at 4°C. The final methylcellulose stock solution was then centrifuged at 15,300 × *g* for 2 hours, and the supernatant was utilized.

PDAC cells were cultured in 24 well plates at densities of 3,000 or 5,000 cells per well. The cells were suspended in 200 μL of culture medium with 1.3% methylcellulose and overlaid on 200 μL of culture medium containing 0.5% agar. The following day, 100 μL of culture medium containing a chemical dissolved in DMSO was added to each well to achieve final DMSO concentrations of 0.1% or 0.2%.

Similarly, PDAC cells were cultured in 48 well plates at densities of 1,000 cells per well. The cells were suspended in 120 μL of culture medium with 1.3% methylcellulose and overlaid on 150 μL of culture medium containing 0.5% agar. The following day, 30 μL of culture medium containing a chemical dissolved in DMSO was added to each well to achieve final DMSO concentrations of 0.1% or 0.2%.

Two weeks post-seeding, 100 μL of fresh medium containing each chemical was added to each well. Three weeks after initiating the culture, spheroids were examined under a microscope, and the number of spheroids of varying sizes was counted.

### Metabolite extraction for comprehensive and target metabolome analyses

To form spheroids, 5,000,000 AsPC-1 or PANC-1 cells were cultured in Cellstar Cell-Repellent Surface culture dishes (Greiner Bio-One GmbH #664970, Frickenhausen, Germany) with media containing 0.65% methylcellulose for 3 days. Subsequently, they were treated with either vehicle, 10 μM RoF, and/or 1 nM Tr. After 24 hours, the spheroids were harvested using 5% mannitol (Fujifilm Wako Pure Chemical #133-00845) and centrifuged at 180 × *g* for 5 min at 4°C, and the supernatants were discarded. The pellets were washed twice with 5% mannitol, resuspended in 1 mL of 5% mannitol, and transferred to Eppendorf tubes. The suspensions were centrifuged at 860 × *g* for 5 min at 4°C, and the supernatants were removed.

To extract metabolites from these cells, the pellets were vortexed and resuspended in 1 mL of methanol (Fujifilm Wako Pure Chemical #138-14521) containing internal standards (25 μM each of methionine sulfone [Alfa Aesar #3A17027, Massachusetts, USA], 2-[N-morpholino]ethanesulfonic acid [MES, Dojindo #341-01622, Kumamoto, Japan], and D-camphor-10-sulfonic acid [CSA, Fujifilm Wako Pure Chemical]), and stored at -20°C.

For comprehensive analysis, 400 µL from the suspension was transferred to a microtube, then 400 µL of chloroform and 200 µL of Milli-Q water were added and mixed well. The solution was centrifuged at 10,000 × *g* for 3 min at 4 °C, and the separated 400 µL aqueous layer was centrifugally filtered through a 5-kDa-cutoff filter (Human Metabolome Technologies #UFC3LCCNB-HMT, Tsuruoka, Japan) to remove proteins. The filtrate was dried using a centrifuge concentrator and reconstituted with 25 µL of Milli-Q water containing reference compounds (200 µmol/L each of 3-aminopyrrolidine [Sigma-Aldrich #404624] and trimesic acid [Fujifilm Wako Pure Chemical #206-03641]) prior to CE-TOFMS analysis (Yugi et al, 2014).

For target metabolome analysis, 400 µL from the suspension was transferred to a microtube, then 400 µL of Milli-Q water was added and mixed well. The solution was centrifuged at 10,000 × *g* for 20 min at 20 °C, and 400 µL of the supernatant was centrifugally filtered through a 5-kDa-cutoff filter (Human Metabolome Technologies #UFC3LCCNB-HMT) to remove proteins. The filtrate was prior to LC-MS/MS analysis.

### Comprehensive metabolome analysis by CE-TOFMS

A quantitative analysis of ionic metabolites in the spheroids were performed using capillary electrophoresis-time of flight mass spectrometer (Agilent Technologies #G6230B). All CE-TOFMS experiments were performed using an Agilent 7100 Capillary Electrophoresis system (Agilent Technologies, Waldbronn, Germany), an Agilent 6230 TOF LC/MS system (Agilent Technologies), an Agilent 1260 series isocratic HPLC pump (Agilent Technologies #G7110B), a G1603A Agilent CE-MS adapter kit (Agilent Technologies #G1603A) and a G1607A Agilent CE-electrospray ionization (ESI)-MS sprayer kit (Agilent Technologies #G1607A). In anionic metabolites analysis, ESI sprayer was replaced with a platinum needle (Agilent Technologies #G7100-60041) instead of initial stainless steel needle (Soga et al, 2009). Other conditions related to CE-ESI-MS sprayer were identical as received. For CE-MS system control and data acquisition, we used Agilent MassHunter software.

### 1) Cationic metabolome analysis

For cationic metabolome analysis, a fused-silica capillary (50 mm i.d. × 100 cm, Molex #TSP050375, Lisle, USA) filled with 1 mol/L formic acid as the electrolyte was used (Soga and Heiger, 2000). A new capillary was flushed with the electrolyte for 20 min, and the capillary was equilibrated for 4 min by flushing with the electrolyte before each run. Sample solution was injected at 5 kPa for 3 sec and a positive voltage of 30 kV was applied. The temperature of the capillary and sample tray was maintained at 20°C and 4°C, respectively. Methanol/water (50% v/v) containing 0.1 mmol/L hexakis(2,2-difluoroethoxy)phosphazene was delivered as sheath liquid at 10 mL/min. ESI-TOFMS was operated in the positive ion mode, and the capillary voltage was set at 4 kV. The flow rate of heated nitrogen gas (heater temperature, 300°C) was maintained at 10 psig. In TOFMS, the fragmentor, skimmer and Oct RF voltages were set at 75, 50 and 125 V, respectively. Automatic recalibration of each acquired spectrum was performed using the masses of reference standards ([^13^C isotopic ion of protonated methanol dimer (2CH_3_OH+H)]^+^, *m/z* 66.06306) and ([hexakis(2,2-difluoroethoxy)phosphazene + H]^+^, *m/z* 622.02896). Exact mass data were acquired at the rate of 1.5 cycles/s over a 50 to 1,000 *m/z* range. Compounds discussed in this paper are listed in Table EV7.

### 2) Anionic metabolome analysis

For anionic metabolome analysis, a COSMO(+) capillary (50 mm i.d. × 105 cm, Nacalai Tesque #07584-44) filled with 50 mmol/L ammonium acetate (pH 8.5) as the electrolyte was used (Soga et al, 2009). Before to the first use, a new capillary was flushed successively with the electrolyte, 50 mmol/L acetic acid (pH 3.4), and then the electrolyte again for 10 min each. Before each run, the capillary was equilibrated by flushing with 50 mmol/L acetic acid (pH 3.4) for 2 min and then with the electrolyte for 5 min. Sample was injected at 5 kPa for 30 sec and a negative voltage of 30 kV was applied. The temperature of the capillary and sample tray was maintained at 20°C and 4°C, respectively. Ammonium acetate (5 mmol/L) in 50% (v/v) methanol/water solution that contained 0.1 mmol/L hexakis(2,2-difluoroethoxy)phosphazene was delivered as sheath liquid at 10 mL/min. ESI-TOFMS was operated in the negative ion mode, and the capillary voltage was set at 3.5 kV. The flow rate of heated nitrogen gas (heater temperature, 300°C) was maintained at 10 psig. In TOFMS, the fragmentor, skimmer, and Oct RF voltages were set at 100, 50, and 200 V, respectively. Automatic recalibration of each acquired spectrum was performed using the masses of reference standards ([^13^C isotopic ion of deprotonated acetate dimer (2CH_3_COOH−H)]^−^, m/z 120.03841) and ([hexakis(2,2-difluoroethoxy)phosphazene + deprotonated acetate(CH_3_COOH−H)]^−^, *m/z* 680.03554). Exact mass data were acquired at the rate of 1.5 cycles/s over a 50 to 1,000 *m/z* range.

### 3) Data analysis of ionic metabolites

The raw data were processed using our proprietary software (MasterHands) (Hirayama et al, 2009; Sugimoto et al, 2010). This software followed the typical processing steps that detected all possible peaks, subtracted baselines, eliminated redundant features (e.g. isotopic, adduct, and fragment peaks), salt and neutral peaks, and noise peaks (e.g. spike peaks), and generated aligned data matrices (Sugimoto et al, 2012). The peaks were identified by matching *m/z* values and normalized migration times of corresponding authentic standard compounds.

### Target metabolome analysis by LC-MS/MS

A quantitative analysis of RF metabolites in the spheroids was performed using LC-triple quadrupole mass spectrometer (Agilent Technologies #G6490A).

The LC system was the Agilent 1290 Infinity HPLC (Agilent Technologies). This system was equipped with automatic degasser (Agilent Technologies #G7122A), binary pump (Agilent Technologies #G4220A), thermostated column compartment (Agilent Technologies #G1316C), autosampler (Agilent Technologies #G4226A) and autosampler cooler (Agilent Technologies #G1330B). Chromatographic separation was performed using an ACQUITY UPLC HSS T3 column (2.1 mm i.d. ×50 mm, 1.8 µm; Waters #186003538, Milford, USA), at a flow rate of 0.3 mL/min and at 45°C. Eluent A consisted of Milli-Q water and eluent B of acetonitrile, both containing 0.1% formic acid and 5 µM medronic acid. Gradient was as follows: 0 min, 0% B; 10 min, 60% B. The injection volume was 1 µL.

The mass spectrometer system was the Agilent 6490 LC/TQ (Agilent Technologies). This system was equipped with Agilent Jet Stream ESI (Agilent Technologies #1958-65138). The MS parameters were set as the following: dry gas temperature 200°C, flow 14 L/min, nebulizer 50 psi, sheath gas temperature 250°C, sheath gas flow 11 L/min, capillary voltage 3500 V, nozzle voltage 1500 V, i-Funnel high pressure RF 90V/low pressure 60V in positive ion mode (Hampel et al, 2012). The ion transitions and optimized parameters for RF metabolite detections are shown in Table EV8. For LC-MS/MS system control and data acquisition, we used Agilent MassHunter software.

RF, FMN, FAD, and RoF were quantified using the internal standard method. Calibration curves for eleven-levels ranging from 2 to 5000 nM were obtained with standard solutions containing the analytes at the respective concentrations spanning the expected ranges in spheroids. Finally, the RF metabolite concentrations were corrected by total protein content.

### RNA-sequence

To form spheroids, 1,000,000 AsPC-1 or PANC-1 cells were cultured in quintuplicate using Nunclon Sphera 60 mm dishes (Thermo Fisher Scientific #174944) with medium containing 0.65% methylcellulose for 3 days. Subsequently, spheroids were treated with either vehicle, 10 μM RoF, and/or 1 nM Tr for 24 hours. Spheroids were collected using PBS, centrifuged at 180 × *g* for 5 min at 4°C, and the supernatants were discarded. The cell pellets were then washed with 1 mL of PBS, transferred to Eppendorf tubes, and centrifuged at 860 × *g* for 5 min at 4°C. After discarding the supernatants, the cell pellets were stored at -80°C until RNA extraction. Total RNA was extracted utilizing RNeasy Mini Kit (Qiagen #74104, Hilden, Germany). To minimize batch differences, five samples of equal RNA quantity were pooled as one representative sample for subsequent analyses. The quality check of RNA samples was performed using Agilent 2100 Bioanalyzer (Agilent, Santa Clara, USA). cDNA library was prepared with TruSeq Standard mRNA Library Prep Kit (Illumina, #20020594). RNA sequencing was performed using NovaSeq6000 with Reagent NovaSeq6000 S4 Reagent Kit v1.5 (Illumina, #20028312). The reads were aligned to human hg38 using STAR (version 2.7.3a) (Dobin et al, 2013). Mapped reads were then counted using HTSEQ (version 2.0.2) (Putri et al, 2022) and the expression of each RNA was converted to transcripts per kilobase million (TPM). For the extraction of differentially expressed genes (DEGs), we used TPM+1 value to minimize the effect of genes whose expression levels were around the background level. We extracted genes whose expressions were up- or down-regulated after the drug treatment (fold change more than 1.3 or less than 0.75). Using the up- and down-regulated gene sets, we undertook connectivity scoring analysis in the JFCR_LinCAGE database, which compared the gene expression changes of query compound with those of anticancer drugs or molecularly targeted agents in the database (Mashima et al, 2015). The gene expression data have been deposited in the Gene Expression Omnibus (accession number GSE274380).

### Statistical analysis

Statistical analyses were conducted using GraphPad Prism 9 (GraphPad Software Inc, San Diego, USA). Threshold for statistical significance was set at *P* < 0.05.

## Acknowledgments

We acknowledge the Sonoshita Laboratory members, Satoshi Ichikawa and Tsuyoshi Konuma for their critical discussions during this study. We also thank Han Hai, Takuya Otsuka, Reo Satoh, Hao Sun, Risa Mizuochi, Madoka Sato, Mika Yamada, Shunsuke Mori, Tomomi Hironaga, Kanna Kondo, Susumu Ishikawa and Katsura Yamaguchi for their technical support. This work was supported by the following grants: Japan Society for the Promotion of Science KAKENHI Grant Numbers JP20K07558, JP22H04922 (AdAMS, to T.O), 20H03524 and 23H02759 (to M.S.), JP23H04946 (to T.S.), Japan Agency for Medical Research and Development grant numbers JP20lm0203001 (to T.O.), 20ck0106548 (to M.S.), JP21zf0127001 (to T.S.), Takeda Science Foundation grant, Japan Foundation for Applied Enzymology, FY21 Support System for the Collaborative Research of Next-Generation Researchers, Hokkaido University, Collaborative Research Grant for the Development of Leadership of Female Researchers, Hokkaido University (to T.O.), Japan Science and Technology Agency, CREST Grant Number JPMJCR2123, Ministry of Education, Culture, Sports, Science and Technology (MEXT) KAKENHI Grant Number JP23H04946 and World Premier International Research Center Initiative (WPI), and Human Biology-Microbiome-Quantum Research Center (Bio2Q) (to T.S.), MEXT, Japan.

## Author contributions

**Takako Ooshio**: Conceptualization; Data curation; Formal analysis; Validation; Investigation; Funding acquisition; Methodology; Writing–original draft; Writing–review and editing. **Yusuke Satoh**: Investigation; Writing–review and editing. **Hui Jiang**: Investigation; Writing–review and editing. **Tongwei Liu**: Investigation; Writing–review and editing. **Kiyonaga Fujii**: Data curation; Investigation; Methodology; Writing–review and editing. **Takamasa Ishikawa**: Data curation; Investigation; Methodology; Writing–review and editing. **Kaori Igarashi**: Investigation; Writing–review and editing. **Kaori Saitoh**: Investigation; Writing–review and editing. **Keiko Kato**: Investigation; Writing–review and editing. **Tetsuo Mashima**: Data curation; Investigation; Methodology; Writing–review and editing. **Shigeki Jin**: Data curation; Investigation; Methodology; Writing–review and editing. **Kotaro Matoba**: Resources; Writing–review and editing. **Matthias Mack**: Resources; Writing–review and editing. **Tsuyoshi Osawa**: Data curation; Writing–review and editing. **Hiroyuki Seimiya**: Methodology; Writing–review and editing. **Tomoyoshi Soga**: Resources; Methodology; Funding acquisition; Writing–review and editing. **Masahiro Sonoshita**: Conceptualization; Resources; Data curation; Investigation; Supervision; Funding acquisition; Methodology; Writing–original draft; Writing–review and editing.

## Ethics statement

All animal studies were conducted using protocols approved by the Hokkaido University Safety Committee on Genetic Recombination Experiments (approval numbers: 2019-007 and 2022-029) and the Hokkaido University Animal Research Committee (approval numbers: 19-0121, 2022-0117, and 2022-0118).

---

**Figure EV1.**
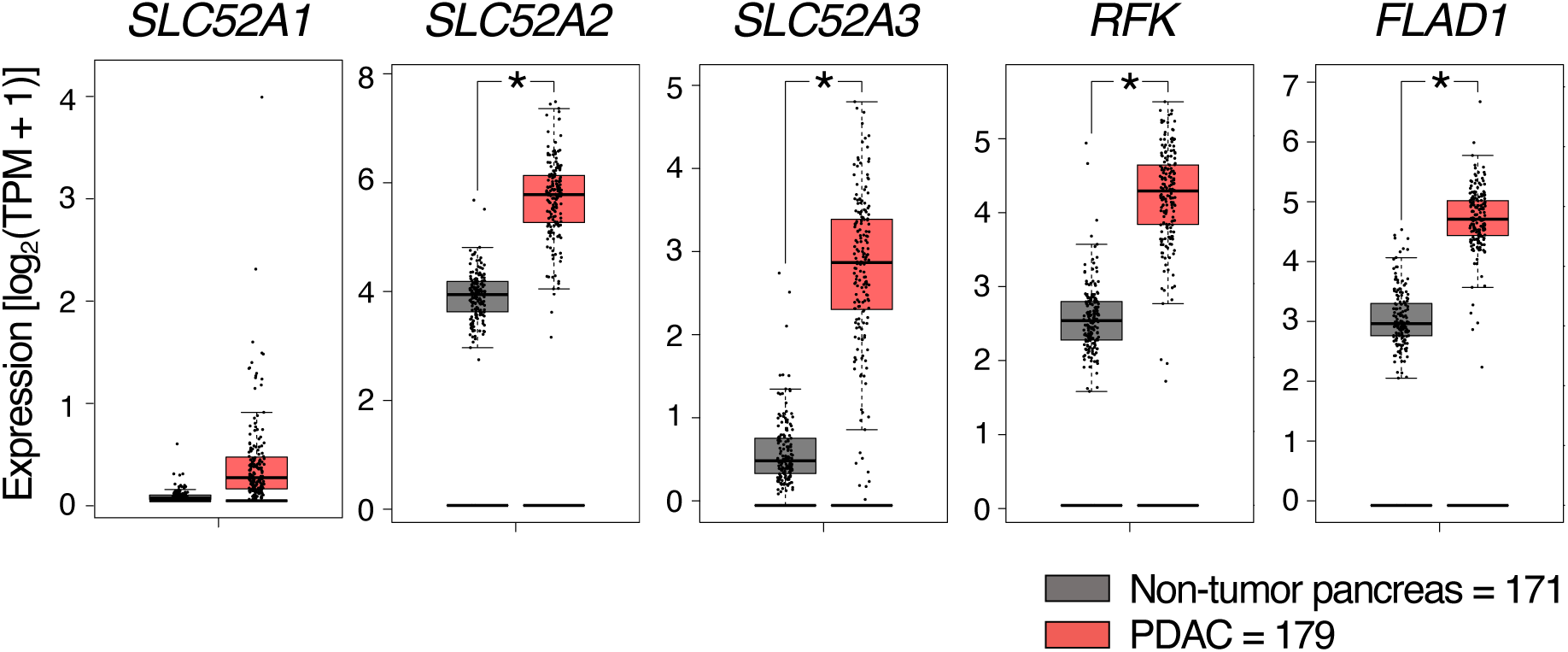
RF metabolism may be activated in PDAC. Expression of RF metabolism-related genes, except *SLC52A1*, was significantly elevated in PDAC compared to non-tumor pancreases. These data were obtained from GEPIA2 (http://gepia2.cancer-pku.cn/#index), which utilizes information from The Cancer Genome Atlas (TCGA) and Genotype-Tissue Expression (GTEx) databases. TMP, transcripts per million. *, *P* < 0.05 in one-way ANOVA.

**Figure EV2.**
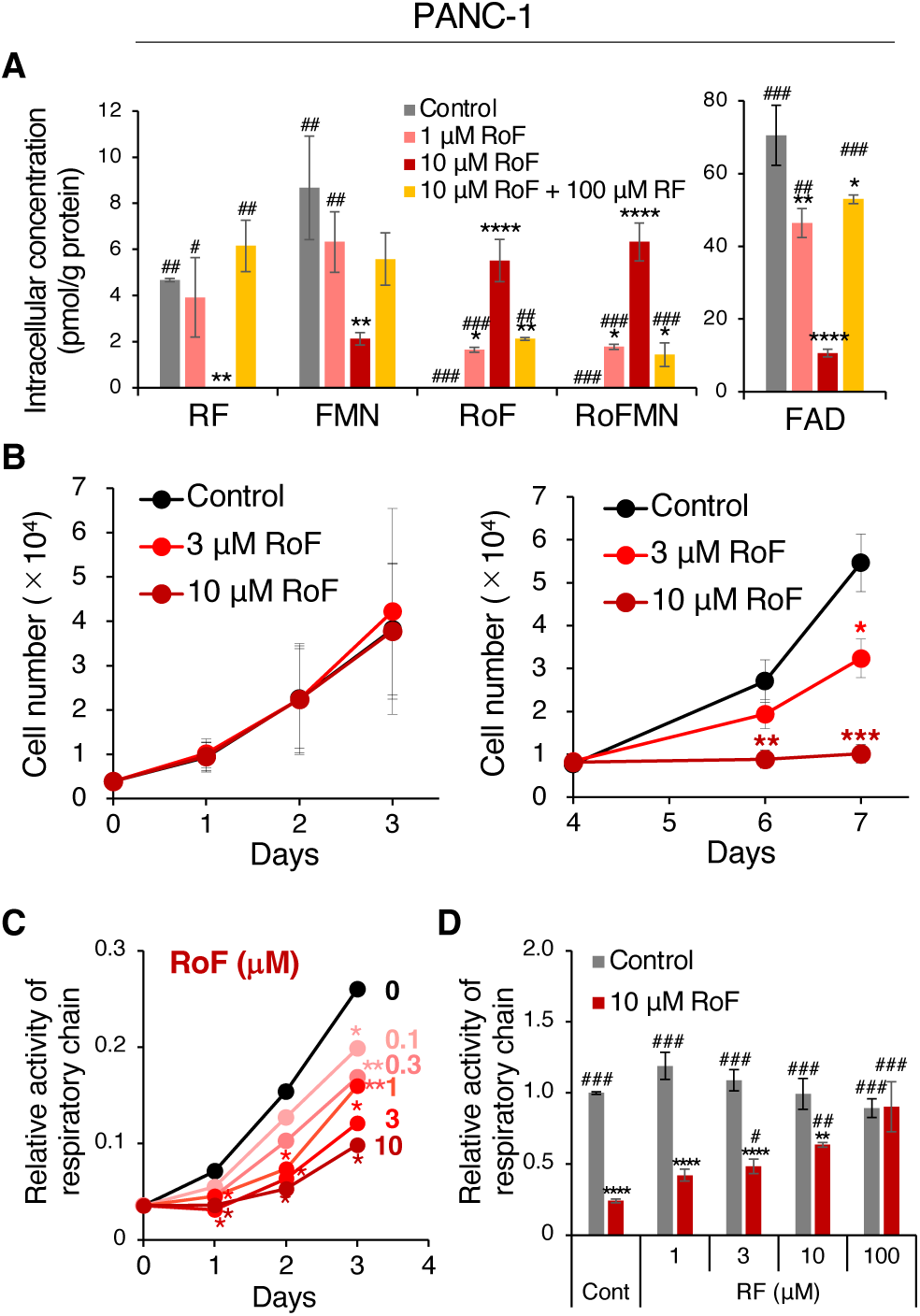
Inhibition of RF metabolism reduces cell growth and mitochondrial respiratory chain activity in PANC-1 cells. (A) RoF treatment decreased intracellular levels of RF and its metabolites by competitively inhibiting RF uptake and metabolism. Metabolite concentrations were quantified 48 hours after treatment with vehicle, RoF, and/or RF. Error bars, SD in technical triplicate. (B) Long-term RoF treatment leads to a reduction in cell numbers. Viable cells were counted using trypan blue exclusion after treatment with vehicle or RoF. (C) RoF inhibits mitochondrial respiratory chain activity in a concentration-dependent manner. Activity was quantified using an MTS assay kit. (D) The inhibition of mitochondrial respiratory chain activity by RoF was reversible with the addition of RF. Respiratory chain activity was reassessed three days post-treatment. *, *P* < 0.05**; \*\***, *P* < 0.005**; \*\*\***, *P* < 0.0005**; \*\*\*\***, *P* < 0.0001 in Dunnett’s test compared to control. ^#^, *P* < 0.05; ^##^, *P* < 0.005; ^##^, *P* < 0.0001 in Dunnett’s test compared to 10 μM RoF.

**Figure EV3.**
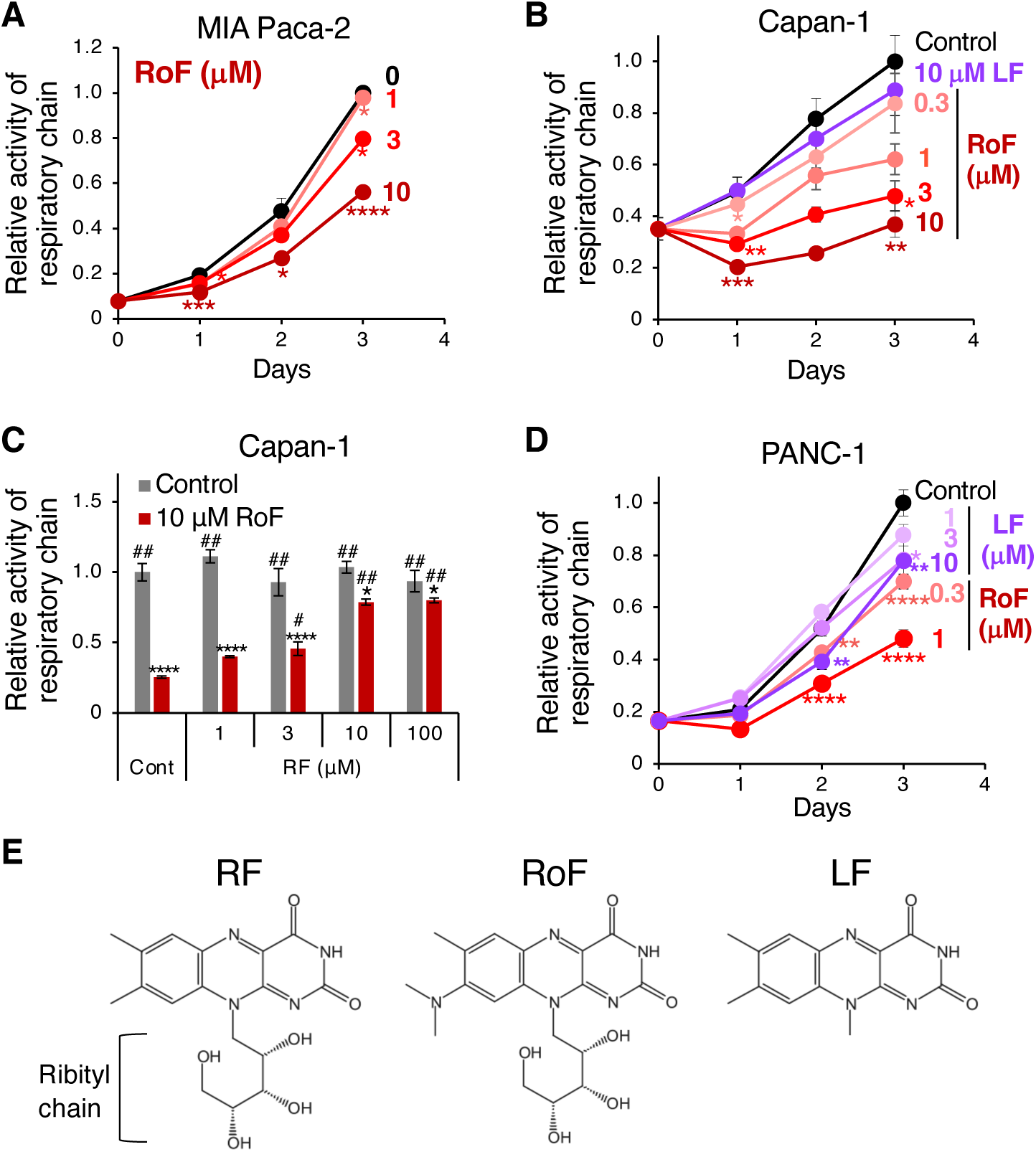
Inhibition of RF metabolism reduced mitochondrial respiratory chain activity in PDAC cells. (A-D) Mitochondrial respiratory chain activity was assessed using MTS assays in MIA PaCa-2 (A), Capan-1 (B and C), and PANC-1 (D) cells following treatment with RoF, lumiflavin (LF), or additional RF. Respiratory chain activity was reassessed three days post-treatment (C). Error bars, SD in technical triplicate. *, *P* < 0.05; **, *P* < 0.01; ***, *P* < 0.005; ****, *P* < 0.001 in Dunnett’s test compared with control. ^#^, *P* < 0.05; ^##^, *P* < 0.0001 in Dunnett’s test compared to 10 μM RoF. (E) Structural comparison of RF, RoF, and LF. LF lacks the ribityl chain, which is the site phosphorylated by RFK.

**Figure EV4.**
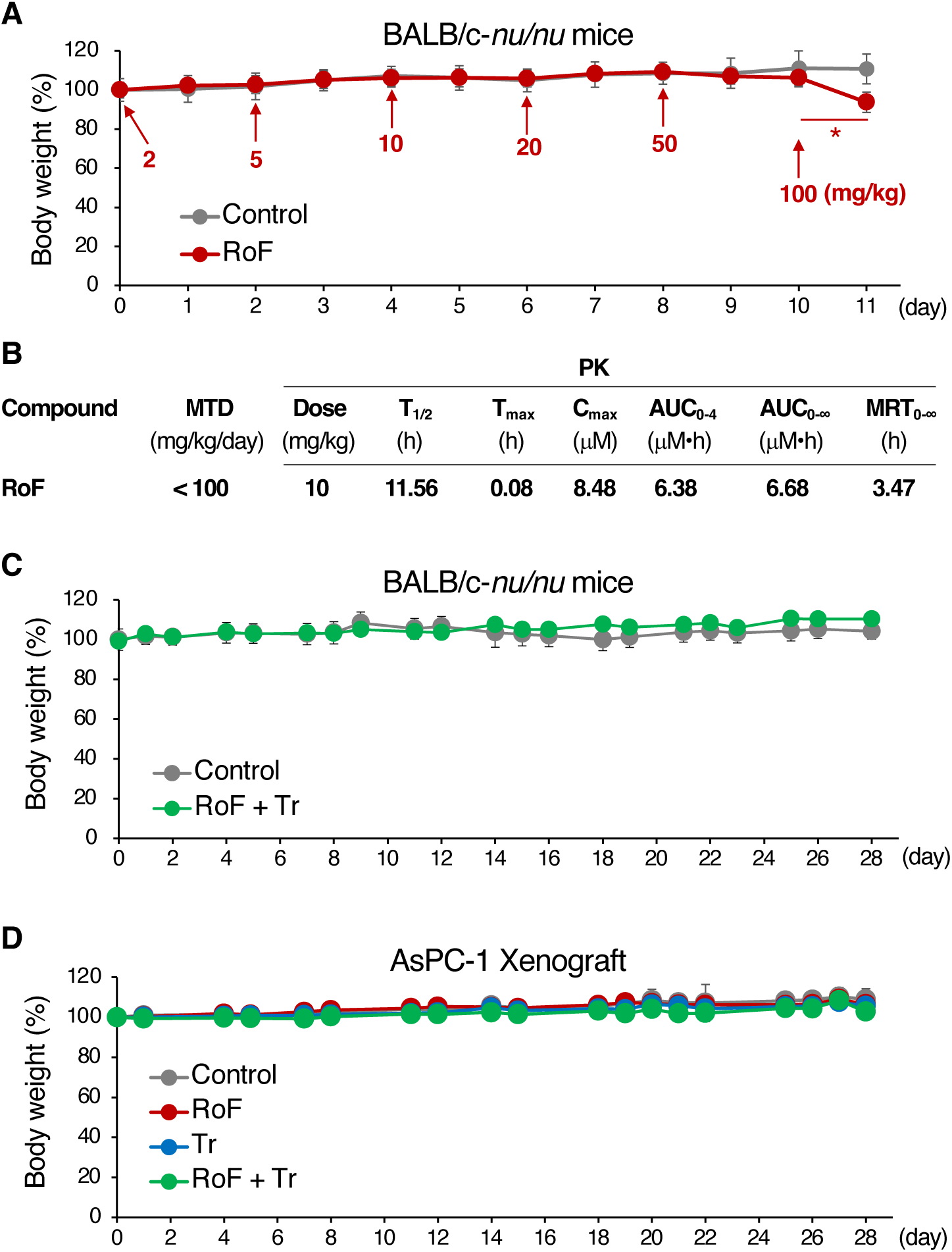
Chemical test in mice. (A) Determining the maximum tolerated dose (MTD) of RoF in mice. Three female BALB/c- *nu*/*nu* mice received an intraperitoneal injection of vehicle (control), while five received increasing doses of RoF. No weight loss or abnormal behavior was observed in mice administered up to 50 mg/kg of RoF. However, significant weight loss occurred the day following administration of 100 mg/kg RoF, resulting in the death of two out of five mice the subsequent day. Therefore, the MTD was determined to be less than 100 mg/kg. Error bars, SD. *, *P* < 0.05 in two-tailed *t*-test. (B) Pharmacokinetic (PK) properties of RoF. Three male ICR mice were dosed intraperitoneally 10 mg/kg RoF. Plasma concentrations of RoF were determined at 0.083, 0.25, 0.5, 1, 2, 4, 8, and 24 hours post-dosing. Error bars, SD. (C) Tolerance of RoF and Tr in mice. Five female BALB/c-*nu*/*nu* mice were administrated 0.2 mg/kg of Tr orally and 10 mg/kg of RoF intraperitoneally, five times per week. Mouse body weight was recorded prior to compound administration, and the weight on Day 0 were set as 100%. Error bars, SD in 5 mice. (D) Tolerance of RoF and Tr in AsPC-1 xenograft mice. Body weight changes were monitored in the same cohort of mice used in the experiment depicted in Figure 3. Control and RoF (n = 7), Tr and RoF + Tr (n = 6). Error bars, SD.

**Figure EV5.**
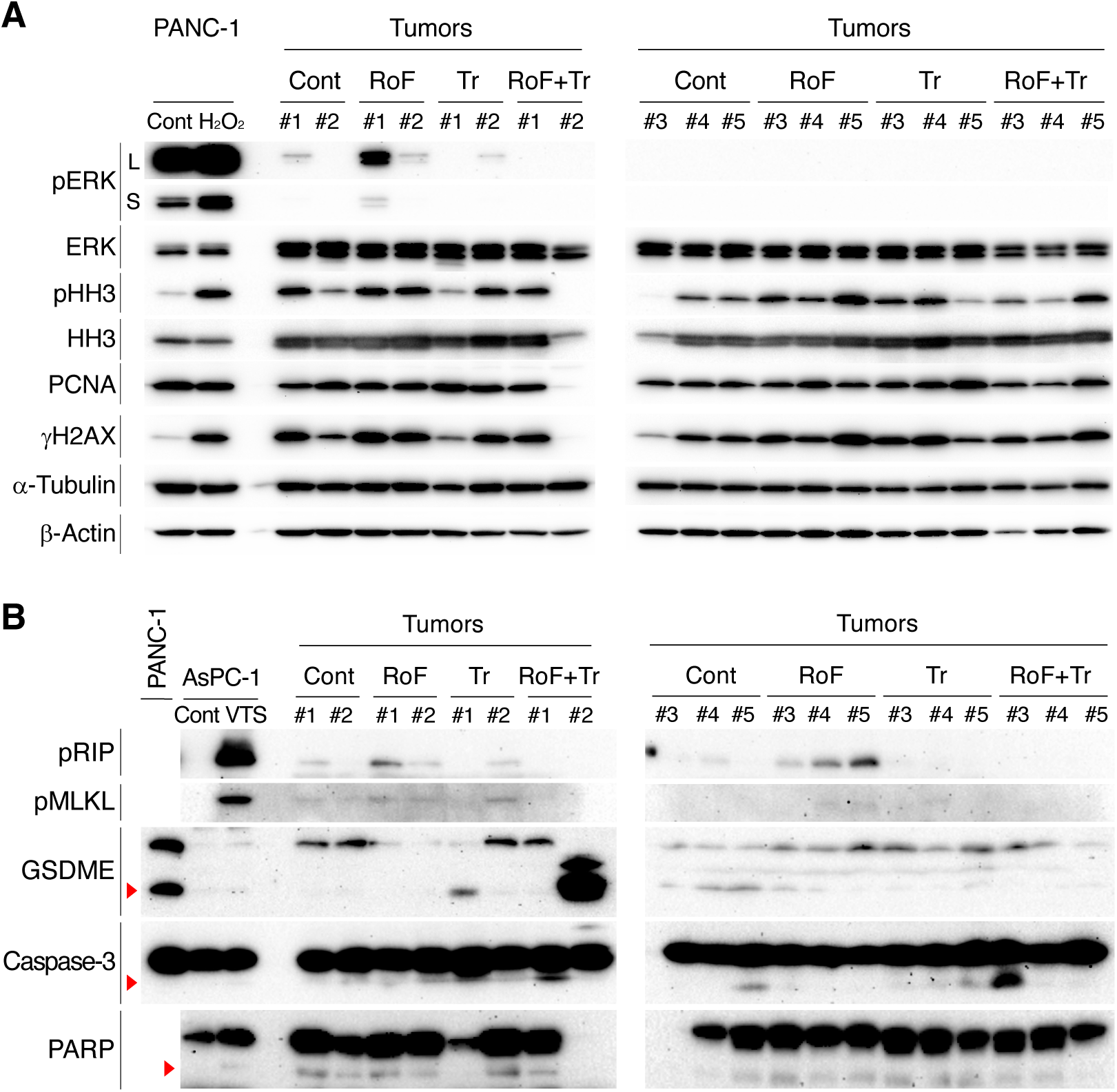
The effects of RoF and/or Tr treatment on the expression of proliferation and cell death markers in PDAC tumors. (A) RoF and/or Tr treatment did not alter the expression of cell proliferation or oxidative stress markers in AsPC-1 xenograft tumors. Thirty μg of protein extracted from the tumors of five mice analyzed in Fig. 3B were loaded per lane. As a positive control for an oxidative stress condition, 23 μg of lysate from PANC-1 cells, which had been treated with either vehicle or 0.5 mM hydrogen peroxide for 1 hour, was also analyzed. The expression of phosphorylated ERK (pERK) was negligible in most tumors and was not noticeably suppressed by Tr treatment. Phosphorylated histone H3 (pHH3) and PCNA were examined as cell proliferation markers, γH2AX as an oxidative stress marker, and α-Tubulin and β-Actin served as internal controls. L, long exposure; S, short exposure. (B) The expression of markers associated with necroptosis, apoptosis, or pyroptosis was not affected by RoF and/or Tr treatment in AsPC-1 xenografts. Sixty μg of proteins extracted from tumors were loaded per lane. For necroptosis controls, AsPC-1 cells were cultured in media containing 20 μM zVAD for 30 min, followed by supplementation with 20 ng/mL TNFα and 100 μM SM-164 for 7 hours (VTS). Thirty-seven μg of tumor protein samples in tumors were loaded per lane. We observed that PANC-1 cells express both the full-length and cleaved forms of GSDME. Forty µg of PANC-1 cell protein was loaded specifically for detecting GSDME and Caspase-3. The expression of phosphorylated RIP (pRIP) and phosphorylated MLKL (pMLKL) as necroptosis markers, cleaved PARP and Caspase-3 as apoptosis markers, and cleaved GSDME as a pyroptosis marker showed no increase following RoF and/or Tr treatment. Red arrowheads, cleaved proteins.

**Figure EV6.**
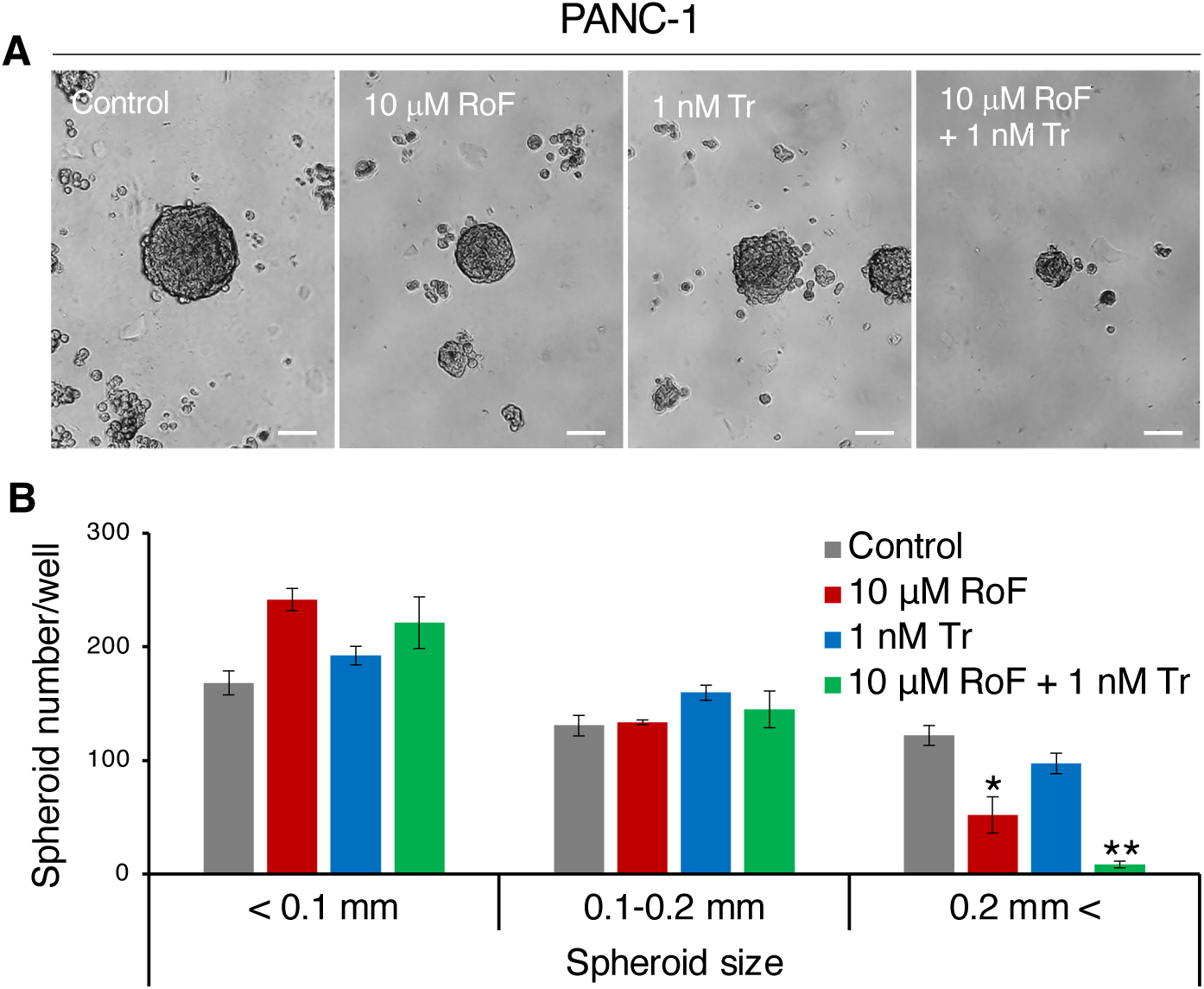
Combined inhibition of RF metabolism and MEK more effectively suppresses spheroid formation in PANC-1 cells than either treatment alone. (A) Three-dimensional culture of PANC-1 cells in methylcellulose media three weeks post-seeding. Scale bars, 0.1 mm. (B) Quantification of spheroids by diameter size, showing a significant reduction in spheroid formation with combined treatment. Error bars, SD in technical triplicate. *, *P* < 0.05; **, *P* < 0.01 in Dunnett’s test compared to control.

**Figure EV7.**
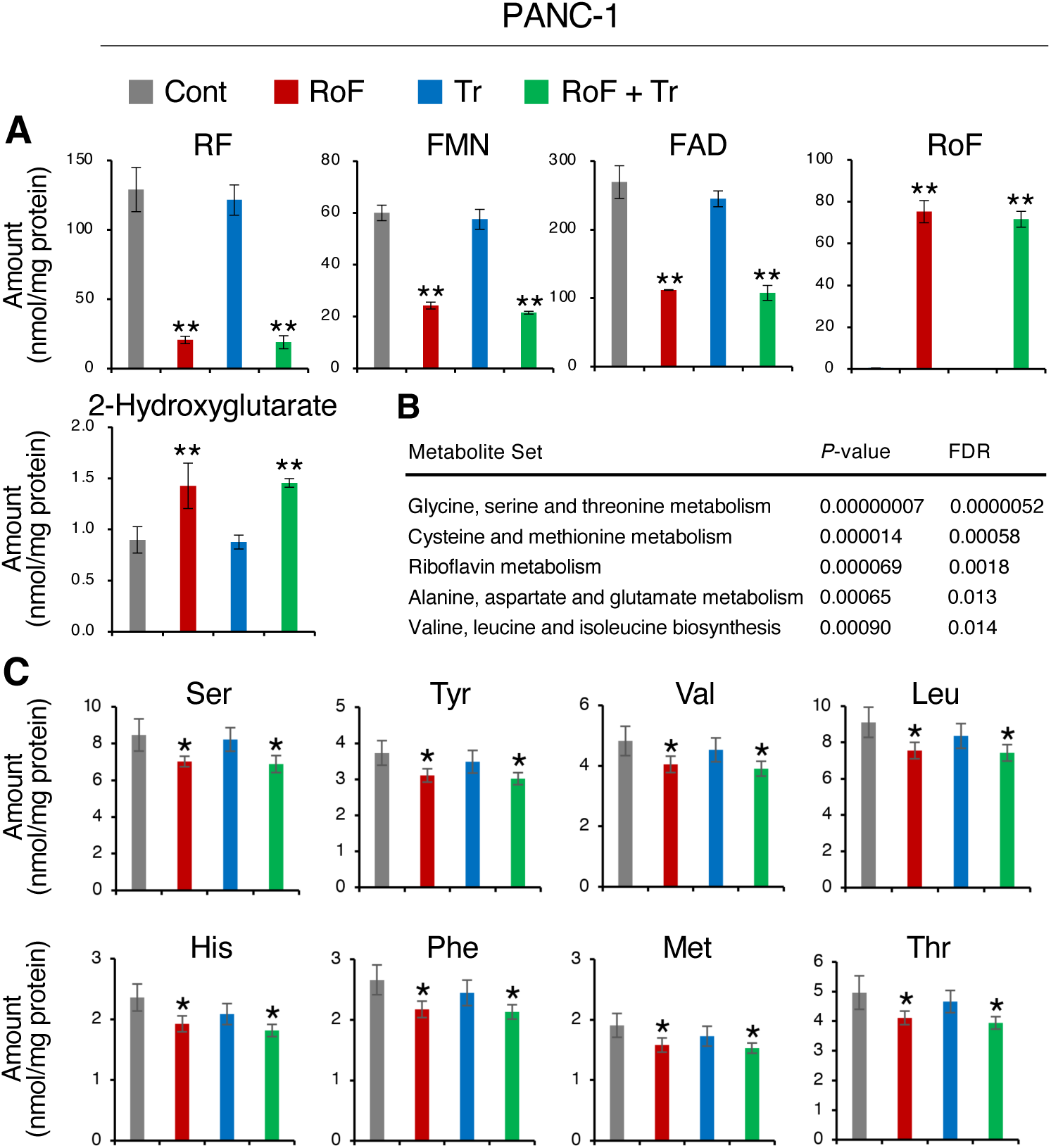
Inhibition of RF metabolism and MEK reduces RF metabolites and several amino acids in PANC-1 cells. (A) RoF and/or Tr treatment led to intracellular uptake of RoF and reduced levels of RF metabolites. (B) Enrichment analysis demonstrated that combined treatment with RoF and Tr significantly reduced levels of amino acids and RF metabolism. Metabolites in cells treated with RoF and Tr, which showed a decrease of 1.2-fold or more compared to control cells, were analyzed using the Metaboanalyst 6.0 (https://www.metaboanalyst.ca/). (C) Treatment with RoF and/or Tr decreased intracellular levels of several amino acids. Error bars, SD in technical quadruplicate. *, *P* < 0.05; **\*\***, *P* < 0.01 in Dunnett’s test compared to control.

**Figure EV8.**
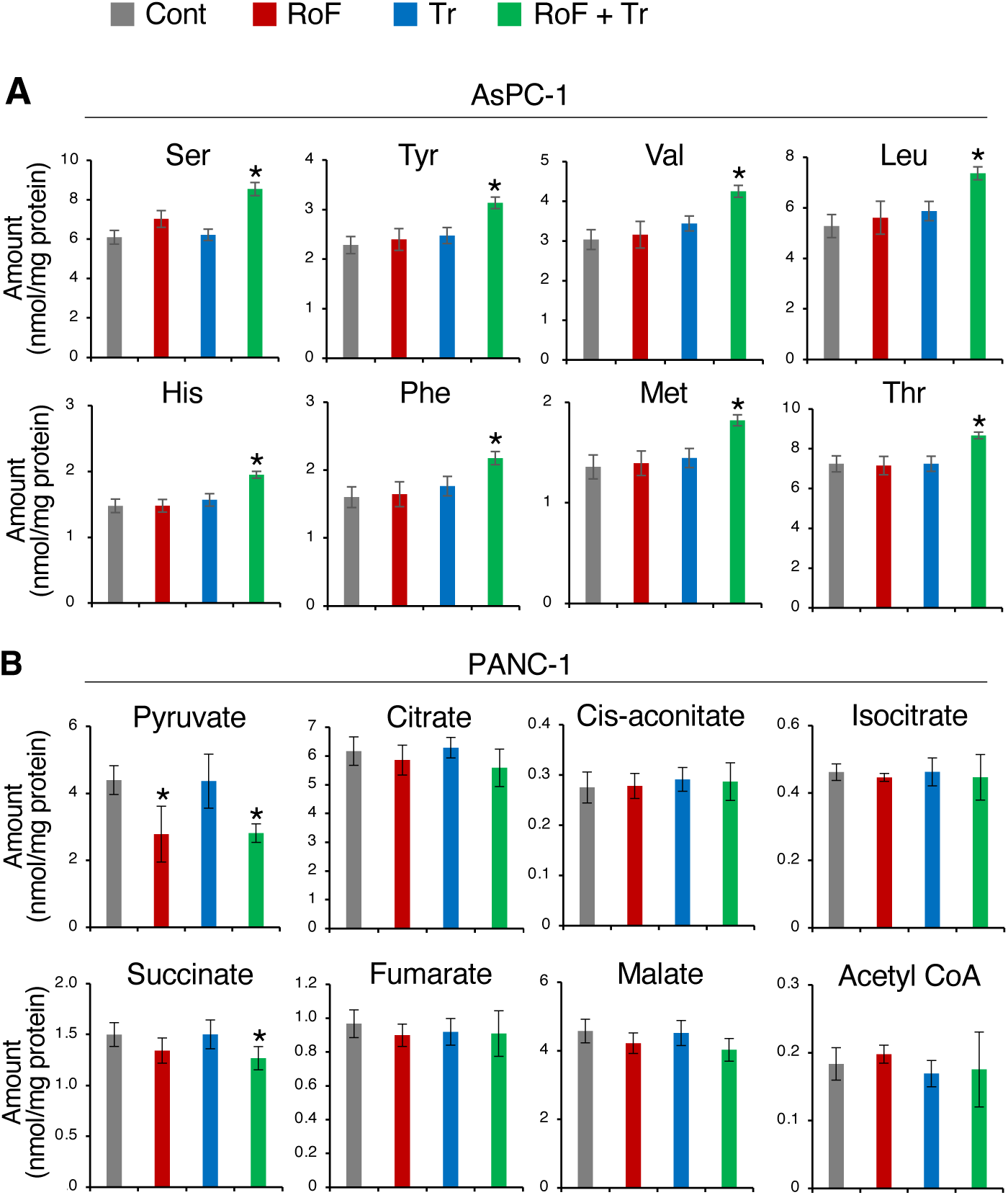
Inhibition of the RF metabolism and MEK increases several amino acids in AsPC-1 cells and decreases some of TCA cycle metabolites in PANC-1 cells. (A) Treatment with combination of RoF and Tr led to an increase in several amino acid levels in AsPC-1 cells. (B) The same combination treatment results in reduced levels of pyruvate and succinate in PANC-1 cells. Error bars, SD in technical quadruplicate. *, *P* < 0.05 in Dunnett’s test compared to control.

**Figure EV9.**
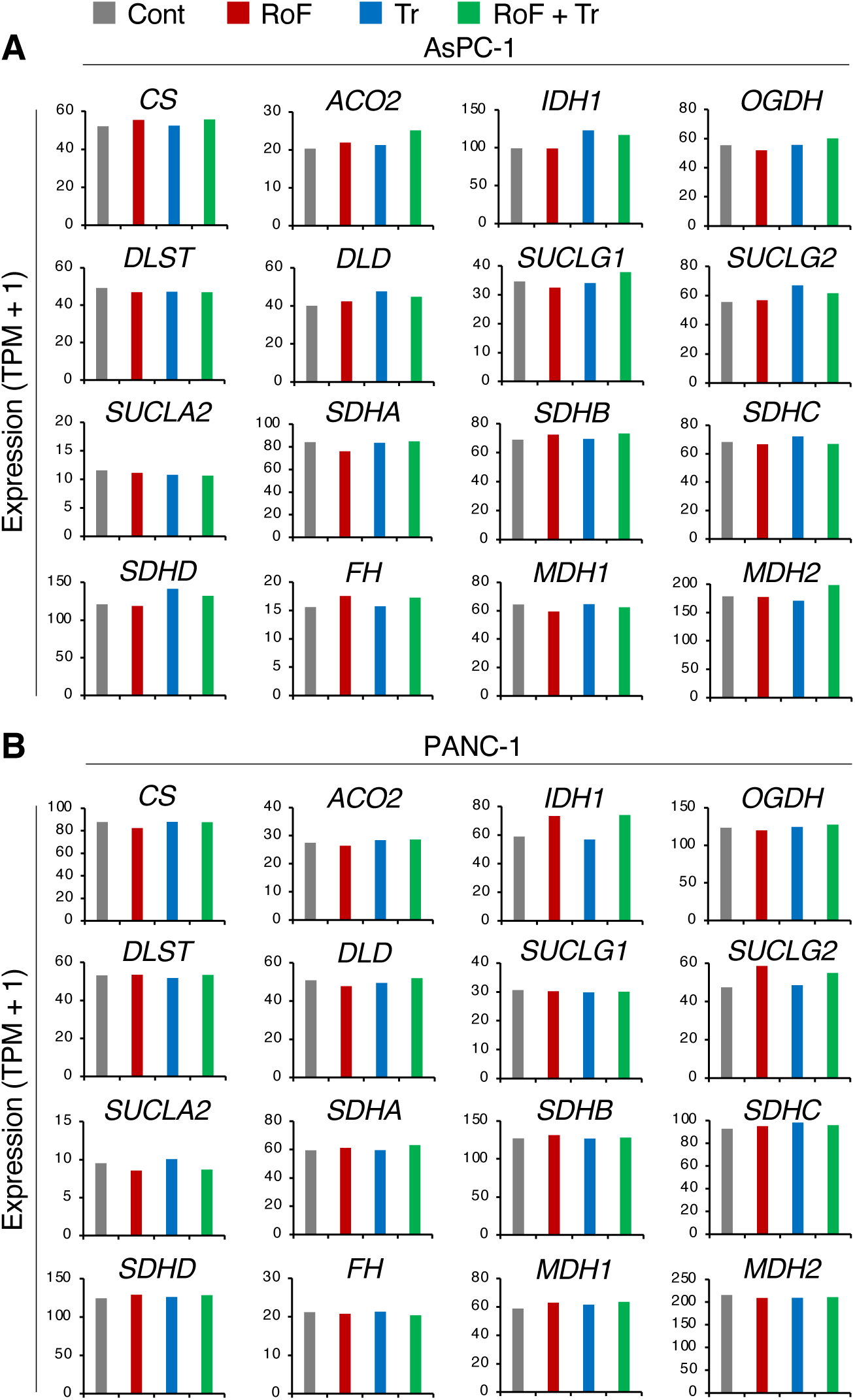
RoF and/or Tr treatment does not decrease the expression of genes related to the TCA cycle. (A and B) Expression of the TCA cycle-related genes based on RNA-sequencing results in AsPC-1 (A) and PANC-1 (B) cells.

**Figure EV10.**
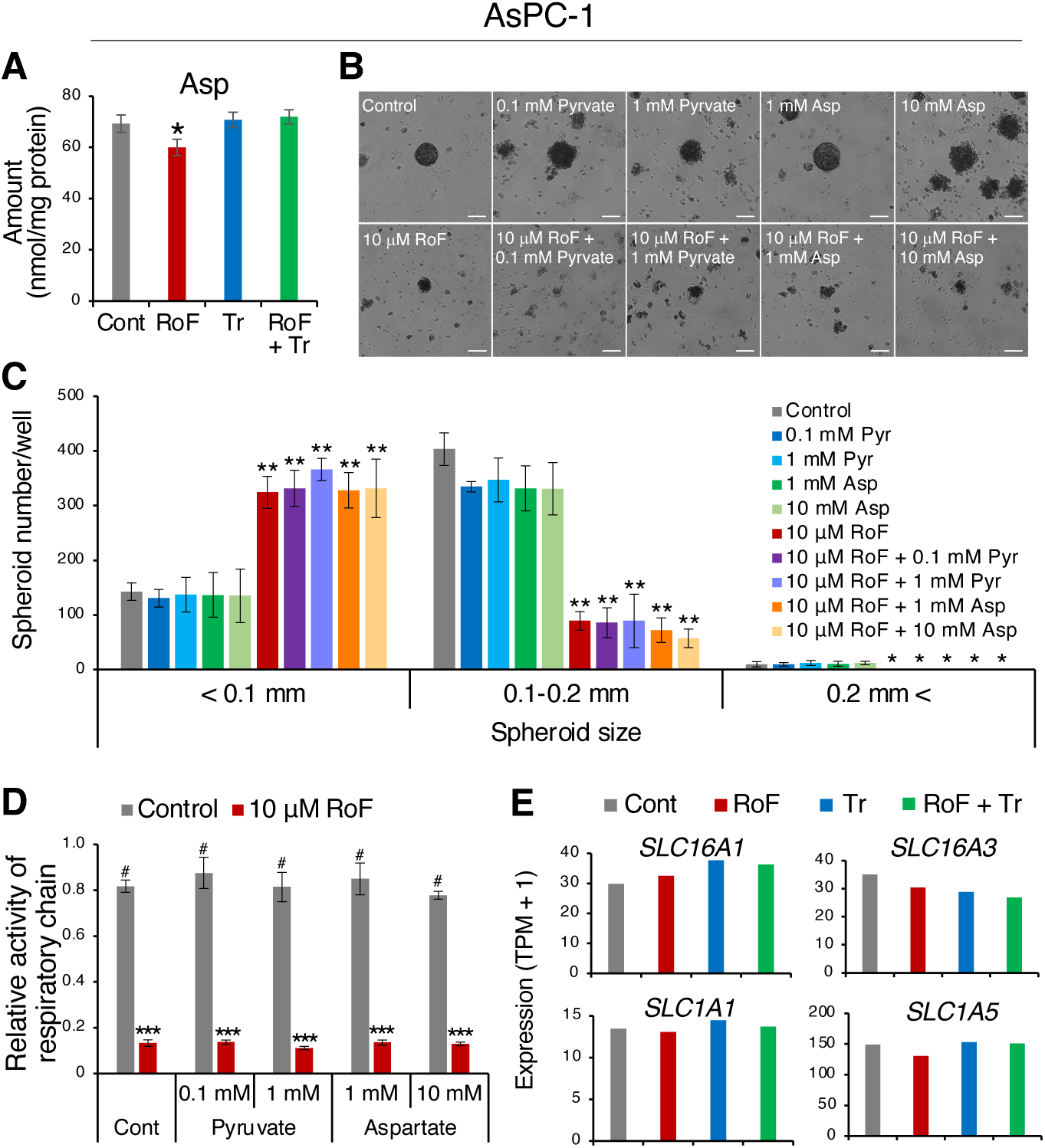
Supplementation of pyruvate or aspartate does not restore the inhibition of spheroid formation by RF metabolism inhibition in AsPC-1 cells. (A) Aspartate level was reduced following RoF treatment. (B) Three-dimensional cultures of AsPC-1 cells in methylcellulose media three weeks post-seeding. Scale bars, 0.1 mm. (C) Quantification of the number of spheroids by diameter size category following treatment with either pyruvate (Pyr) or aspartate (Asp). (D) Supplementation with either Pyr or Asp did not restore respiratory chain activity inhibited by RoF treatment. (E) RNA-sequencing data showed expression of Asp (*SLC16A1* and *SLC16A3*) and Pyr (*ALC1A1* and *ALC1A5*) transporters following treatment. Error bars, SD in technical triplicate. *, *P* < 0.05; **, *P* < 0.01; ***, *P* < 0.005 in Dunnett’s test compared to control. ^#^, *P* < 0.005 in Dunnett’s test compared to 10 μM

**Figure EV11.**
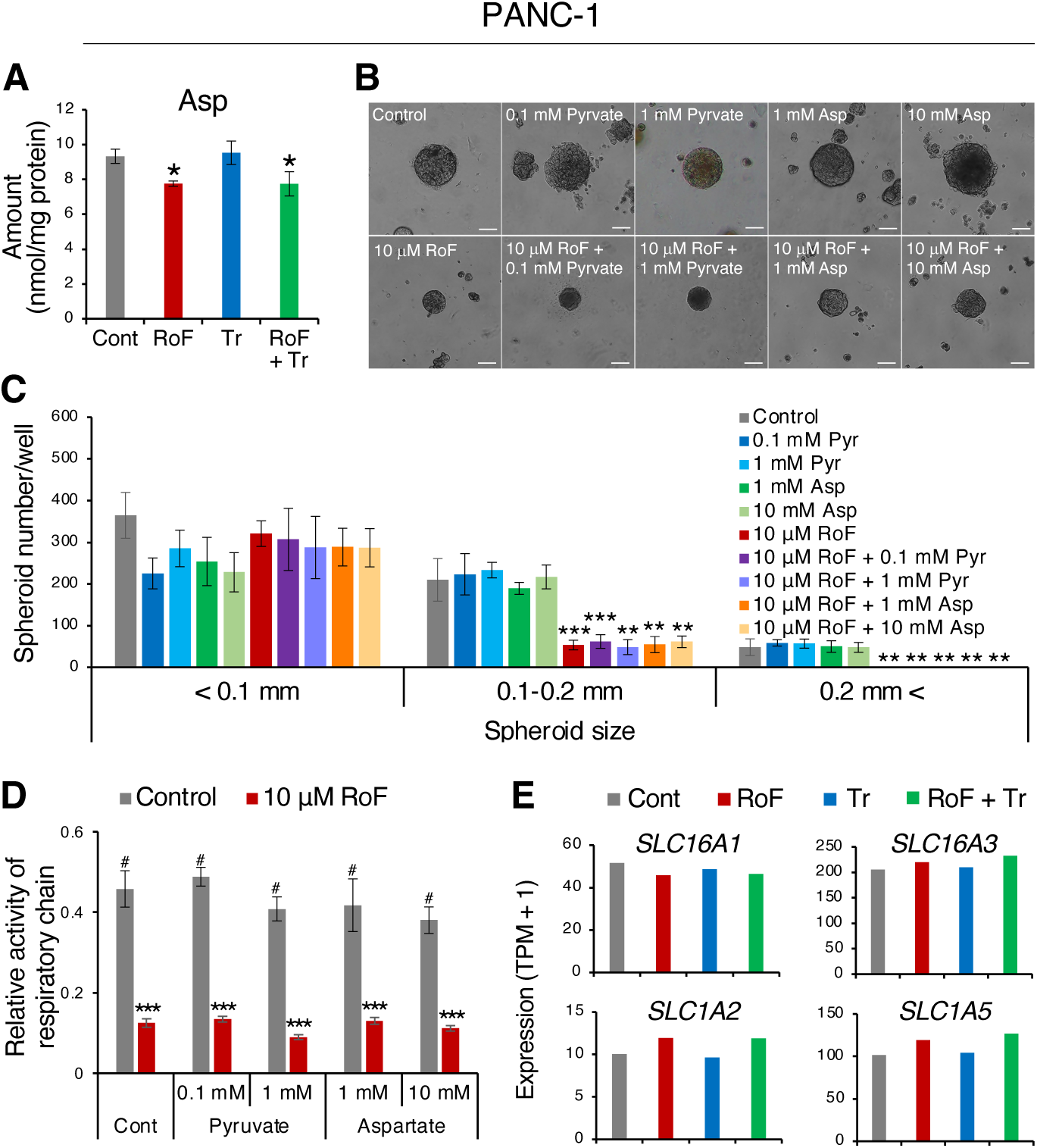
Supplementation of pyruvate or aspartate does not restore the inhibition of spheroid formation by RF metabolism inhibition in PANC-1 cells. (A) Aspartate level was decreased following RoF or RoF + Tr treatment. (B) Three-dimensional cultures of PANC-1 cells in methylcellulose media three weeks post-seeding. Scale bars, 0.1 mm. (C) Quantification of the number of spheroids by diameter size category following treatment with either Pyr or Asp. (D) Supplementation of Pyr or Asp did not restore respiratory chain activity inhibited by RoF treatment. (E) RNA-sequencing data showed expression of Asp (*SLC16A1* and *SLC16A3*) or Pyr (*ALC1A2* and *ALC1A5*) transporters. Error bars, SD in technical triplicate. *, *P* < 0.05; **, *P* < 0.01; ***, *P* < 0.005 in Dunnett’s test compared to control. ^#^, *P* < 0.005 in Dunnett’s test compared to 10 μM RoF.

**Figure EV12.**
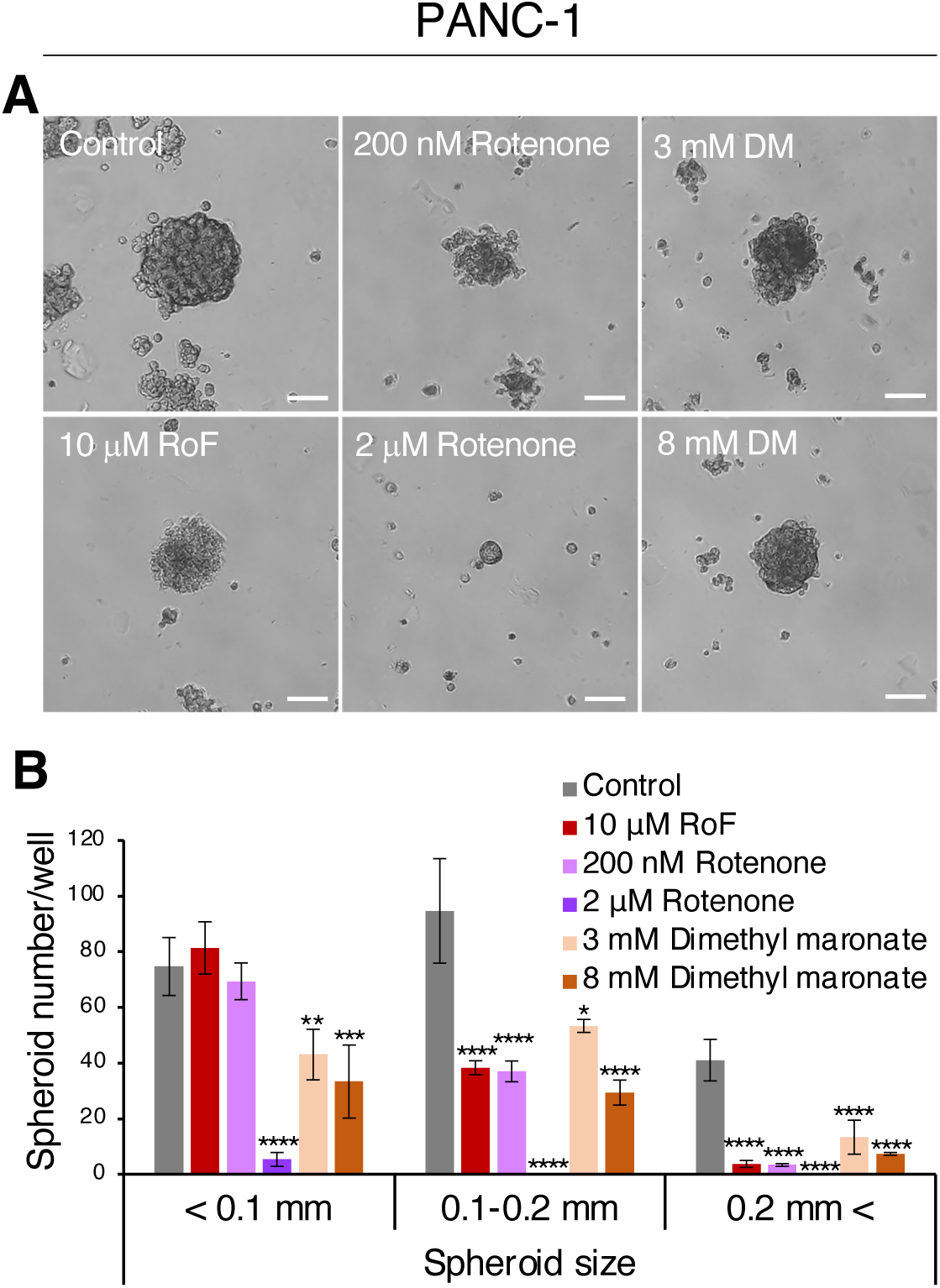
Inhibition of mitochondrial respiratory chain reduces spheroid formation in PANC-1 cells. (A) Three-dimensional cultures of PANC-1 cells in methylcellulose media treated with RoF, rotenone, or dimethyl malonate (DM) three weeks post-seeding. Scale bars, 0.1 mm. (B) Quantification of spheroids by diameter size, showing a reduction in spheroid formation following treatment with respiratory chain inhibitors. Error bars, SD in technical triplicate. *, *P* < 0.05; **, *P* < 0.01; ***, *P* < 0.005; ****, *P* < 0.0001 in Dunnett’s test compared to control.

**Table EV1.** Metabolites reduced by 1.2-fold or more in PANC-1 cells following treatment with RoF and/or Tr.

| Downregulated metabolites in PANC-1 cells |  |  |  |  |  |  |  |  |  |  |  |
| --- | --- | --- | --- | --- | --- | --- | --- | --- | --- | --- | --- |
| KEGG ID | Compound name | FC (R/C) | P | KEGG ID | Compound name | FC (T/C) | P | KEGG ID | Compound name | FC (RT/C) | P |
| C00295 | Orotate | 0.00 | 0.001 | C00581 | Guanidinoacetate | 0.00 | 0.391 | C00581 | Guanidinoacetate | 0.00 | 0.391 |
| C00255 | RF | 0.16 | 0.001 | C05123 | Isethionate | 0.34 | 0.235 | C00295 | Orotate | 0.00 | 0.001 |
| C08261 | Azelate | 0.24 | 0.132 | C02727 | N-epsilon-Acetyllysine | 0.45 | 0.269 | C00255 | RF | 0.15 | < 0.001 |
| C00061 | FMN | 0.40 | < 0.001 | C00455 | NMN | 0.51 | 0.313 | C10172 | Proline betaine | 0.27 | 0.149 |
| C00016 | FAD | 0.42 | 0.001 | C06772 | Diethanolamine | 0.52 | 0.183 | C00061 | FMN | 0.36 | < 0.001 |
| C00262 | Hypoxanthine | 0.49 | 0.148 | C02997 | N-Acetylhistidine | 0.58 | 0.418 | C00016 | FAD | 0.40 | < 0.001 |
| C00294 | Inosine | 0.50 | 0.117 | C00661 | G3P | 0.59 | 0.447 | C02997 | N-Acetylhistidine | 0.51 | 0.330 |
| C00438 | Carbamoylaspartate | 0.57 | 0.013 | C00857 | Deamido-NAD+ | 0.62 | 0.477 | C00455 | NMN | 0.54 | 0.348 |
| C00022 | Pyruvate | 0.63 | 0.035 | C00160 | Glycolate | 0.65 | 0.513 | C00294 | Inosine | 0.55 | 0.144 |
| C01081 | Thiamine monophosphate | 0.63 | 0.020 | C03451 | S-Lactoylglutathione | 0.65 | 0.243 | C00438 | Carbamoylaspartate | 0.60 | 0.016 |
| C01585 | Hexanoate | 0.68 | 0.288 | C00750 | Spermine | 0.69 | 0.283 | C01081 | Thiamine monophosphate | 0.60 | 0.012 |
| C06772 | Diethanolamine | 0.69 | 0.403 | C00262 | Hypoxanthine | 0.70 | 0.335 | C00131 | dATP | 0.61 | 0.153 |
| C03451 | S-Lactoylglutathione | 0.71 | 0.390 | C00637 | Indole-3-acetaldehyde | 0.70 | 0.047 | C00250 | Pyridoxal | 0.61 | 0.041 |
| C03413 | N1,N12-Diacetylspermine | 0.72 | 0.013 | C00131 | dATP | 0.71 | 0.309 | C00022 | Pyruvate | 0.64 | 0.003 |
| - | Ophthalmate | 0.73 | 0.378 | C00250 | Pyridoxal | 0.75 | 0.169 | C00212 | Adenosine | 0.68 | 0.127 |
| C00006 | NADP+ | 0.73 | 0.007 | C00096 | GDP-mannose | 0.75 | 0.173 | C00366 | Urate | 0.71 | 0.075 |
| C00250 | Pyridoxal | 0.76 | 0.150 | C00705 | dCDP | 0.76 | 0.087 | C00074 | PEP | 0.71 | 0.070 |
| C00197 | 3PG | 0.77 | 0.022 | C01585 | Hexanoate | 0.77 | 0.482 | C00051 | Glutathione(red) | 0.71 | 0.584 |
| C00212 | Adenosine | 0.79 | 0.265 | C02037 | Gly-Gly | 0.80 | 0.524 | C00006 | NADP+ | 0.73 | 0.101 |
| C02291 | Cystathionine | 0.79 | 0.152 | C00383 | Malonate | 0.81 | 0.099 | C00197 | 3PG | 0.74 | 0.015 |
| C00096 | GDP-mannose | 0.79 | 0.038 | C00147 | Adenine | 0.81 | 0.129 | C00383 | Malonate | 0.75 | 0.062 |
| C00051 | Glutathione(red) | 0.79 | 0.729 | C00119 | PRPP | 0.82 | 0.476 | C02727 | N-epsilon-Acetyllysine | 0.76 | 0.621 |
| C00074 | PEP | 0.80 | 0.138 | C01089 | 3-Hydroxybutyrate | 0.82 | 0.086 | C00135 | His | 0.77 | 0.017 |
| C00064 | Gln | 0.80 | 0.032 |  |  |  |  | C00262 | Hypoxanthine | 0.78 | 0.665 |
| C02356 | 2AB | 0.80 | 0.048 |  |  |  |  | C00064 | Gln | 0.79 | 0.025 |
| C00637 | Indole-3-acetaldehyde | 0.81 | 0.084 |  |  |  |  | C01073 | N-Acetyl-beta-alanine | 0.79 | 0.110 |
| C00041 | Ala | 0.81 | 0.090 |  |  |  |  | C00307 | CDP-choline | 0.79 | 0.063 |
| C00135 | His | 0.82 | 0.035 |  |  |  |  | C00188 | Thr | 0.79 | 0.045 |
| C00079 | Phe | 0.82 | 0.032 |  |  |  |  | C00152 | Asn | 0.80 | 0.040 |
| C02571 | o-Acetylcarnitine | 0.82 | 0.272 |  |  |  |  | C01585 | Hexanoate | 0.80 | 0.535 |
| C00153 | Nicotinamide | 0.82 | 0.388 |  |  |  |  | C02356 | 2AB | 0.80 | 0.045 |
| C00750 | Spermine | 0.82 | 0.323 |  |  |  |  | C01005 | O-Phosphoserine | 0.80 | 0.363 |
| C00055 | CMP | 0.83 | 0.346 |  |  |  |  | C02291 | Cystathionine | 0.80 | 0.168 |
| C00152 | Asn | 0.83 | 0.064 |  |  |  |  | C00079 | Phe | 0.80 | 0.025 |
| C01152 | 3-Methylhistidine | 0.83 | 0.088 |  |  |  |  | - | N1,N8-Diacetylspermidine | 0.80 | 0.022 |
| C00188 | Thr | 0.83 | 0.073 |  |  |  |  | C00073 | Met | 0.80 | 0.038 |
| C00123 | Leu | 0.83 | 0.039 |  |  |  |  | C00041 | Ala | 0.80 | 0.083 |
| C00065 | Ser | 0.83 | 0.059 |  |  |  |  | C00078 | Trp | 0.80 | 0.033 |
| C00073 | Met | 0.83 | 0.059 |  |  |  |  | C00082 | Tyr | 0.81 | 0.027 |
| C00049 | Asp | 0.83 | 0.004 |  |  |  |  | C00183 | Val | 0.81 | 0.038 |
| C00082 | Tyr | 0.83 | 0.042 |  |  |  |  | C00063 | CTP | 0.81 | 0.126 |
| C01026 | N,N-Dimethylglycine | 0.83 | 0.277 |  |  |  |  | C00065 | Ser | 0.81 | 0.044 |
|  |  |  |  |  |  |  |  | C00123 | Leu | 0.82 | 0.030 |
|  |  |  |  |  |  |  |  | C01152 | 3-Methylhistidine | 0.82 | 0.074 |
|  |  |  |  |  |  |  |  | C01157 | Hydroxyproline | 0.82 | 0.062 |
|  |  |  |  |  |  |  |  | C00096 | GDP-mannose | 0.82 | 0.049 |
|  |  |  |  |  |  |  |  | C00327 | Citrulline | 0.83 | 0.073 |
|  |  |  |  |  |  |  |  | C00300 | Creatine | 0.83 | 0.194 |
|  |  |  |  |  |  |  |  | C00719 | Betaine | 0.83 | 0.040 |
|  |  |  |  |  |  |  |  | C00049 | Asp | 0.83 | 0.021 |
|  |  |  |  |  |  |  |  | C00407 | Ile | 0.83 | 0.058 |
|  |  |  |  |  |  |  |  | C03413 | N1,N12-Diacetylspermine | 0.83 | 0.337 |

FC, fold change; R, RoF treatment; C, vehicle treatment (control); T, Tr treatment; RT, combination treatment of RoF and Tr

**Table EV2.**
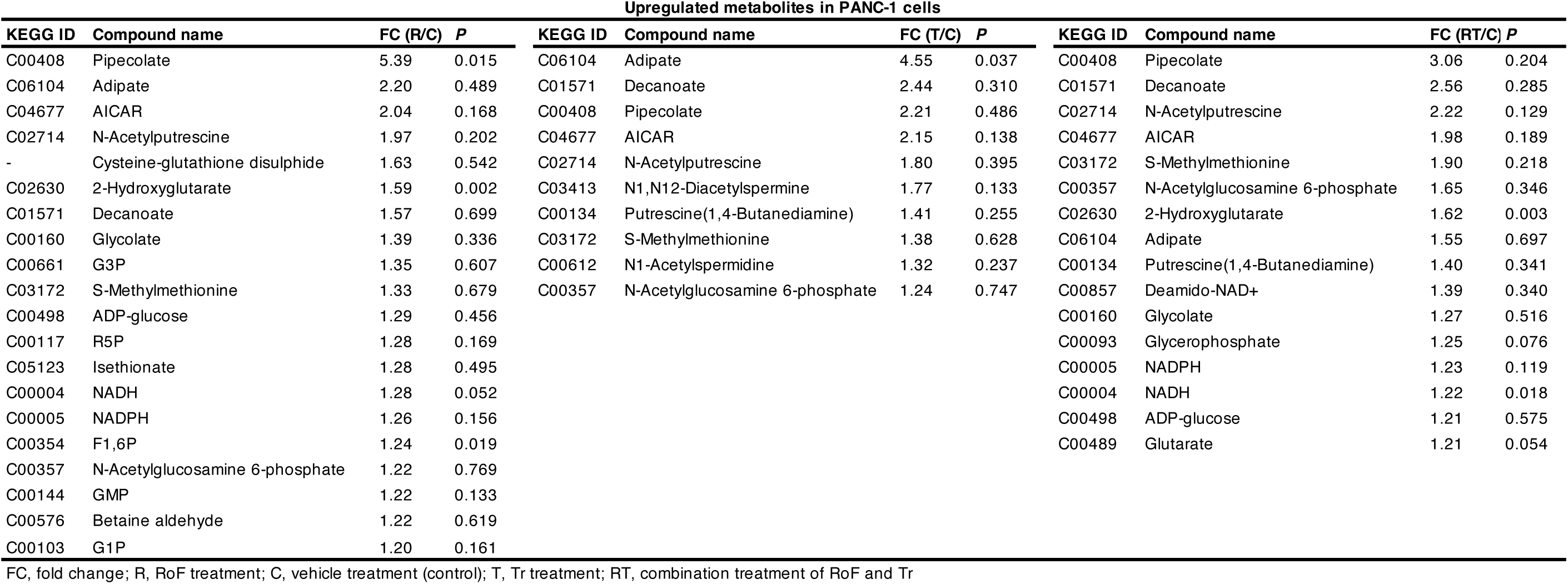
Metabolites elevated by 1.2-fold or more in PANC-1 cells following treatment with RoF and/or Tr.

**Table EV3.** Comparative analysis of RF-related metabolites and mRNA levels in AsPC-1 and PANC-1 cells.

|  | LC-MS analysis (nmol/g protein) |  |  | RNA-seq (TPM+1) |  |  |  |  |
| --- | --- | --- | --- | --- | --- | --- | --- | --- |
|  | RF | FMN | FAD | *SLC52A1* | *SLC52A2* | *SLC52A3* | *RFK* | *FLAD1* |
| AsPC-1 | 1042 ± 91 | 22 ± 4 | 233 ± 14 | 1 | 105 | 17 | 7 | 18 |
| PANC-1 | 129 ± 16 | 60 ± 3 | 269 ± 24 | 2 | 35 | 4 | 17 | 30 |
| *P* | 0.00019 | < 0.0001 | 0.047 | – | – | – | – | – |

**Table EV4.** JFCR_LinCAGE analysis results in AsPC-1 cells.

| AsPC-1 cells |  |  |  |  |  |  |  |  |  |  |  |
| --- | --- | --- | --- | --- | --- | --- | --- | --- | --- | --- | --- |
| Tr |  |  |  |  |  | RoF + Tr |  |  |  |  |  |
| Rank | Sample ID | Compound | Connectivity | Up score | Down score | Rank | Sample ID | Compound | Connectivity | Up score | Down score |
| 1 | C086 | U-0126 | 1.000 | 0.409 | -0.210 | 1 | C085 | AG1024 | 1.000 | 0.168 | -0.176 |
| 2 | C036 | Sorafenib | 0.995 | 0.377 | -0.239 | 2 | C004 | A23187 | 0.992 | 0.145 | -0.196 |
| 3 | C039 | Sunitinib | 0.930 | 0.280 | -0.296 | 3 | C071 | Amrubicin | 0.985 | 0.190 | -0.149 |
| 4 | C121 | Dabrafenib | 0.876 | 0.316 | -0.227 | 4 | C059 | Bortezomib | 0.932 | 0.145 | -0.176 |
| 5 | C114 | Erlotinib30 | 0.874 | 0.299 | -0.243 | 5 | C015 | 5-FU | 0.851 | 0.142 | -0.150 |
| 6 | C113 | Erlotinib0.6 | 0.870 | 0.319 | -0.220 | 6 | C099 | Regorafenib | 0.825 | 0.151 | -0.133 |
| 7 | C037 | Sorafenib | 0.859 | 0.334 | -0.198 | 7 | C010 | Geldanamycin | 0.801 | 0.129 | -0.147 |
| 8 | C124 | Afatinib | 0.857 | 0.291 | -0.240 | 8 | C009 | MG-132 | 0.749 | 0.114 | -0.144 |
| 9 | C099 | Regorafenib | 0.853 | 0.325 | -0.203 | 9 | C121 | Dabrafenib | 0.744 | 0.134 | -0.123 |
| 10 | C116 | Gefitinib30 | 0.840 | 0.339 | -0.181 | 10 | C018 | Mitomycin C | 0.737 | 0.109 | -0.144 |

**Table EV5.** Fly stocks for mutant or RNAi.

| Description | Fly gene | Fly gene ID | Fly genotype | RRID |
| --- | --- | --- | --- | --- |
| *Rfk* -/+ | *Rfk* | CG2846 | *w*[1118]; *P*{*w*[+*mC*]=*EP*}*CG2846*[*G4442*] | BDSC_30081 |
| *Rift* RNAi #1 | *Rift* | CG11576 | *y*[1] *v*[1]; *P*{*y*[+*t*7.7] *v*[+*t*1.8]=*TRiP.HM05210*}*attP2/TM3, Sb*[1] | BDSC_29531 |
| *Rift* RNAi #2 | *Rift* | CG11576 | *Rift* RNAi | NIG_11576R-3 |
| *Rfk* RNAi #1 | *Rfk* | CG2846 | *y*[1] *sc*[*] *v*[1] *sev*[21]; *P*{*y*[+*t*7.7] *v*[+*t*1.8]=*TRiP.HMC04709*}*attP40* | BDSC_30084 |
| *Rfk* RNAi #2 | *Rfk* | CG2846 | *P*{*KK107342*}*VIE-260B* | VDRC_100266 |
| *Flad1* RNAi #1 | *Flad1* | CG4407 | *y*[1] *sc*[*] *v*[1] *sev*[21]; *P*{*y*[+*t*7.7] *v*[+*t*1.8]=*TRiP.HMS02772*}*attP40* | BDSC_44055 |
| *Flad1* RNAi #2 | *Flad1* | CG4407 | *FADS* RNAi | NIG_4407R-4 |
| *Flad2* RNAi #1 | *Flad2* | CG16848 | *y*[1] *v*[1]; *P*{*y*[+*t*7.7] *v*[+*t*1.8]=*TRiP.HMJ23521*}*attP40* | BDSC_61938 |
| *Flad2* RNAi #2 | *Flad2* | CG16848 | *Flad2* RNAi | NIG_16848R-1 |

RRID, Research Resource Identifiers

**Table EV6.** Media to culture human PDAC cell lines.

| Cell line | RRID | Medium | FBS | Penicillin/streptomycin |
| --- | --- | --- | --- | --- |
| AsPC-1 | CVCL_0152 | Roswell Park Memorial Institute 1640 | 10% | 1% |
| PANC-1 | CVCL_0480 |  |  |  |
| Capan-1 | CVCL_0237 | Iscove's Modified Dulbecco's Medium | 20% | 1% |
| MIA PaCa-2 | CVCL_0428 | Dulbecco's minimal essential medium | 10% | 1% |

FBS, fetal bovine serum. Final concentrations of FBS and penicillin/streptomycin are indicated.

**Table EV7.**
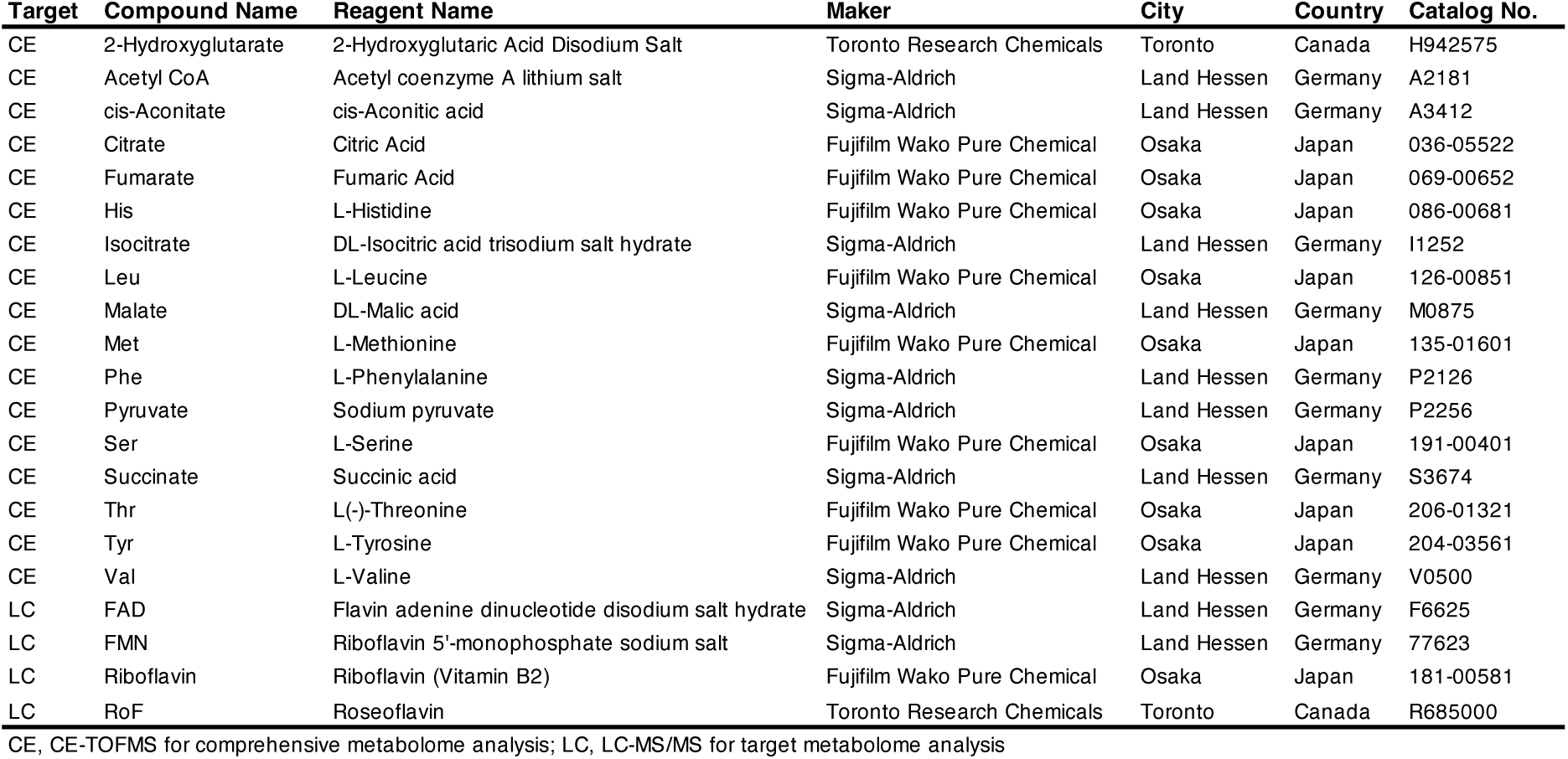
Compound list for comprehensive and target metabolome analysis.

| Target | Compound Name | Reagent Name | Maker | City | Country | Catalog No. |
| --- | --- | --- | --- | --- | --- | --- |
| CE | 2-Hydroxyglutarate | 2-Hydroxyglutaric Acid Disodium Salt | Toronto Research Chemicals | Toronto | Canada | H942575 |
| CE | Acetyl CoA | Acetyl coenzyme A lithium salt | Sigma-Aldrich | Land Hessen | Germany | A2181 |
| CE | cis-Aconitate | cis-Aconitic acid | Sigma-Aldrich | Land Hessen | Germany | A3412 |
| CE | Citrate | Citric Acid | Fujifilm Wako Pure Chemical | Osaka | Japan | 036-05522 |
| CE | Fumarate | Fumaric Acid | Fujifilm Wako Pure Chemical | Osaka | Japan | 069-00652 |
| CE | His | L-Histidine | Fujifilm Wako Pure Chemical | Osaka | Japan | 086-00681 |
| CE | Isocitrate | DL-Isocitric acid trisodium salt hydrate | Sigma-Aldrich | Land Hessen | Germany | I1252 |
| CE | Leu | L-Leucine | Fujifilm Wako Pure Chemical | Osaka | Japan | 126-00851 |
| CE | Malate | DL-Malic acid | Sigma-Aldrich | Land Hessen | Germany | M0875 |
| CE | Met | L-Methionine | Fujifilm Wako Pure Chemical | Osaka | Japan | 135-01601 |
| CE | Phe | L-Phenylalanine | Sigma-Aldrich | Land Hessen | Germany | P2126 |
| CE | Pyruvate | Sodium pyruvate | Sigma-Aldrich | Land Hessen | Germany | P2256 |
| CE | Ser | L-Serine | Fujifilm Wako Pure Chemical | Osaka | Japan | 191-00401 |
| CE | Succinate | Succinic acid | Sigma-Aldrich | Land Hessen | Germany | S3674 |
| CE | Thr | L(-)-Threonine | Fujifilm Wako Pure Chemical | Osaka | Japan | 206-01321 |
| CE | Tyr | L-Tyrosine | Fujifilm Wako Pure Chemical | Osaka | Japan | 204-03561 |
| CE | Val | L-Valine | Sigma-Aldrich | Land Hessen | Germany | V0500 |
| LC | FAD | Flavin adenine dinucleotide disodium salt hydrate | Sigma-Aldrich | Land Hessen | Germany | F6625 |
| LC | FMN | Riboflavin 5'-monophosphate sodium salt | Sigma-Aldrich | Land Hessen | Germany | 77623 |
| LC | Riboflavin | Riboflavin (Vitamin B2) | Fujifilm Wako Pure Chemical | Osaka | Japan | 181-00581 |
| LC | RoF | Roseoflavin | Toronto Research Chemicals | Toronto | Canada | R685000 |

CE, CE-TOFMS for comprehensive metabolome analysis; LC, LC-MS/MS for target metabolome analysis

**Table EV8.** MS/MS ion transitions monitored for target metabolome analysis.

| Compound | Precursor ion | Product ion | Dwell time (msec) | Frag (V) | CE (V) | Cell Acc (V) | Polarity |
| --- | --- | --- | --- | --- | --- | --- | --- |
| RF | 377 | 243 | 200 | 380 | 24 | 5 | Positive |
| FMN | 457 | 243 | 200 | 380 | 30 | 5 | Positive |
| RoF | 406 | 272 | 200 | 380 | 28 | 5 | Positive |
| FAD | 786 | 348 | 200 | 380 | 20 | 5 | Positive |
| CSA (IS) | 233 | 123 | 200 | 380 | 17 | 5 | Positive |

IS, internal standard; CE, collision energy

